# Genome-wide selection on transposable elements in maize

**DOI:** 10.1101/2025.09.16.676665

**Authors:** Beibei Liu, Manisha Munasinghe, Regina A. Fairbanks, Candice N. Hirsch, Jeffrey Ross-Ibarra

## Abstract

While most evolutionary research has focused on single nucleotide polymorphisms (SNPs), transposable elements (TEs) represent a major but understudied source of mutations that can influence organismal fitness. Previous studies on TEs often overlook the mechanisms and rates of transposition, rely on short-read sequencing that limits TE detection, or focus on small genomes such as *Arabidopsis* or *Drosophila*. In this study, we leveraged high-quality, long-read genome assemblies from 26 maize inbreds to investigate natural selection on TEs. We developed a novel and interpretable method, Φ_*SFS*_, which incorporates TE age and improves resolution for detecting selection. Using this approach, we identified key factors influencing selection on TEs: (1) the distance to the nearest gene, (2) the pre-insertion DNA methylation level at the insertion site, and (3) intrinsic TE characteristics, including copy number and expression level. This work represents the first application of long-read genome assemblies to study TE selection in a major crop species with a typical plant genome size. Our Φ_*SFS*_method offers a broadly applicable framework for detecting selection on TEs, and the factors uncovered provide new insights into the evolutionary dynamics and trade-offs between TEs and host genes.

## Introduction

Genetic variation is the fundamental driver of adaptation. While most insights into the genetic basis of adaptation have come from studies of single nucleotide polymorphisms (SNPs) due to their abundance and ease of detection, the effects of individual SNPs are typically small. Transposable elements (TEs), or “jumping genes,” represent another major source of genetic variation. First discovered in maize by Barbara McClintock (<u>McClintock 1950</u>), TEs — selfish elements that replicate via RNA (Class I) or DNA (Class II) intermediates — have since been found across the tree of life. While single nucleotide changes generally have limited impacts on the genome, TEs can impact the genome in a myriad of ways. For example, TEs can create (Studer et al. 2011) or destroy (Kobayashi et al. 2004) regulatory sequence, move or duplicate genes (Xiao et al. 2008), knock out genes (Bhattacharyya et al. 1990), or provide the substrate for epigenetic regulation of key phenotypes (Ong-Abdullah *et al*. 2015). As such, TEs often exhibit much larger impacts on phenotype than SNPs. Even in small genomes, TEs are responsible for a substantial portion of segregating variation in expression in rice (Li et al. 2024) and in flies TE mutations alone can provide sufficient genetic variation in quantitative traits for selection to act on (Torkamanzehi et al. 1992).

While the effects of TE insertions imply they should impact fitness, genome-wide characterization of selection on TEs has been limited. Efforts to look at many TEs have predominantly focused on compact genomes, often evaluate only a subset of TEs, or rely on genomic approaches with poor genotyping accuracy. And while some work has identified individual insertions that are beneficial (Studer *et al*. 2011; Hof *et al*. 2016; Kobayashi *et al*. 2004), or context-dependent (Butelli *et al*. 2012), and TEs have been shown to contribute to quantitative traits like flowering time (Yang *et al*. 2013), most theoretical (Munasinghe *et al*. 2023b) and empirical (Hollister and Gaut 2009; Song and Boissinot 2007) work has treated TEs as unambiguously deleterious. For example, new insertions in rice are notably lacking from functional regions such as exons (Naito *et al*. 2006) and negative associations between TE copy number and fitness are taken as evidence of widespread selection against TEs (Stitzer *et al*. 2023). Nonetheless, our knowledge of the selective consequences of TEs is still limited by the lack of genome-wide polymorphism data.

Polymorphic TE data across populations are essential for understanding selection on TEs. However, accurately identifying TEs using short-read sequencing is challenging due to their repetitive nature. In most plant genomes, TEs make up the vast majority of sequence, and short-read-based methods are incapable of highly accurate genotyping (Menard *et al*. 2025). The advent of long-read genome assembly, however, now enables more accurate detection of TE polymorphisms, and recent efforts have used these technologies to identify TEs genomewide in a number of plant species (Li *et al*. 2024; Wei *et al*. 2022; Munasinghe *et al*. 2023a).

Given accurate genotyping of TEs polymorphisms, analysis of the TE site frequency spectrum (SFS) is a potentially powerful approach to infer selection (Nielsen 2005). Comparison of the frequency distribution of TEs to neutral expectations has been used in several small-scale studies, almost universally identifying an excess of low frequency insertions (Hollister and Gaut 2009;

Neafsey et al. 2004). While an excess of low frequency mutations is a hallmark of either negative or stabilizing selection (Nielsen 2005; Charlesworth 2013), the theory underlying such inference assumes variants arise at a constant rate, an assumption that does not hold for TEs. Transposition rates are highly variable and can increase in response to genomic stresses or environmental stimuli (O’Neill et al. 1998; Lisch 2009; Hénault et al. 2020; Makarevitch et al. 2015). Recent bursts of transposition would produce an excess of young, low-frequency insertions compared to constant-rate models, mimicking the SFS signature of negative selection. Similarly, older bursts followed by a decrease in transposition rate would lead to an excess of intermediate frequency variants. This confounding effect makes it challenging to disentangle the influence of selection from transposition. To address the confounding effects of transposable element (TE) bursts, Horvath *et al*. (2022) developed an age-adjusted site frequency spectrum for detecting selection on TEs. By matching TEs with SNPs of the same age, Horvath *et al*. (2022) showed via simulation that the expected age-adjusted frequency of neutral TEs matched that of SNPs, and was robust both to variation in the rate of accumulation of variants and population size change. While this approach has been successfully applied to detect selection in TE polymorphism data (Jiang et al. 2024; Horvath *et al*. 2024), to date these analyses have relied on short-read genotyping and have only been done in relatively small plant genomes.

In this study, we leveraged high-quality long-read genome assemblies from 26 maize inbred lines to investigate TE insertion dynamics and the factors influencing their selection patterns. We developed a new statistic Φ_sfs_ to quantify selection on TEs by comparing the SFS of TEs to an age-matched set of genome-wide SNPs. Building on our previous comparisons of age-adjusted allele frequency (Horvath et al. 2022), Φ_sfs_ incorporates TE age and offers more intuitive interpretation. Applying Φ_sfs_ to our maize TE data, we show convincing evidence of selection against TEs near genes, likely due to their impacts on regulatory sequence and therefore gene expression. We also quantify the impact of intrinsic TE features such as family copy number, length, and expression on TE evolution. Finally, based on our results, we argue that the consequences of TEs might be better thought of through the lens of stabilizing selection on quantitative traits rather than one of unambiguously deleterious mutations.

## Materials and Methods

### Polymorphic TE and SNP Datasets

In Munasinghe *et al*. (2023a), base-pair–resolved pairwise alignments were generated between B73 and each of the remaining 25 inbred founder lines of the Nested Association Mapping (NAM) population (Hufford et al. 2021; McMullen et al. 2009). From the resulting gVCF (genomic Variant Call Format) files, structural variants (SVs) were identified. SVs were defined as genomic regions greater than 50 bp in one genotype that were absent in the corresponding region of the other genotype (Figure S1A). Each SV was further classified based on its overlap with transposable element (TE) annotations into one of four categories: ‘No TE SV’, ‘Incomplete TE SV’, ‘TE = SV’, and ‘TE Within SV’ (Figure S1B). To obtain TE polymorphic data across the NAM founder lines, we combined orthologoous ‘TE = SV’ across the 25 pairwise alignments. Each corresponding SV was assigned a unique structural haplotype identifier (see GitHub repository: https://github.com/beibeiliu1008/TE_Evo). Specifically, we grouped all SVs by chromosome and applied this process chromosome by chromosome. We started by taking the posi-2 Liu *et al*. tion, size, and category of all SVs. The first SV in the set is treated as our ‘reference SV’. We create 50 bp buffers around the start and end position (25 bp to left and right) of our reference SV. We then go through every other SV in the set. An SV is grouped with the reference SV if (1) its start position falls within the reference start buffer zone, (2) its end position falls within the reference end buffer zone, (3) the size of the SV is within 50bp of the reference size, and (4) the category matches the category of the reference SV (Figure S2). Based on these criteria, we linked 300,272 “TE==SV” across 25 NAM lines (data available on Github). We use GATK (McKenna *et al*. 2010) to merge gVCFs and call single nucleotide polymorphisms (SNPs) between B73 and each of the remaining 25 NAM lines. We exclude sites with missing data, non-TE indels, and nonbiallelic SNPs were excluded.

### Age estimation of TEs and SNPs

The age estimation of TEs and SNPs was performed using RELATE (Speidel *et al*. 2019), which reconstructs genealogical trees along the genome. To prepare the input files for RELATE, we converted the above TE and SNP polymorphic data (VCF files) into .haps files using `RelateFileFormats --mode ConvertFromVcf` (scripts available on GitHub: https://myersgroup.github.io/relate/input_data.html, https://github.com/beibeiliu1008/TE_Evo). We made an ancestral reference genome by replacing the B73V5 genome with the SNPs between B73 and *Zea diploperrenis*. We also generated a masked version of the reference genome, retaining only the start positions (1 bp) of the identified TE insertions, the positions of invariant SNPs, and the positions of non-missing biallelic SNPs, while all other regions were masked. We used a mutation rate of 3.3 × 10^−8^ per base pair per generation (Clark *et al*. 2005) and an effective population size of 300,000 (Beissinger *et al*. 2016). While the insertion rate of TEs per bp is likely lower than the per-nucleotide mutation rate, the vast majority of polymorphisms in the combined data file are SNPs, so the above mutation rate is reasonable for the combined data. The .haps files of TEs and SNPs were combined for each chromosome, and each resulting .haps and .sams file was used as input to estimate the age of each SNP and TE by executing `RelateParallel.sh` (https://github.com/beibeiliu1008/TE_Evo). Each chromosome’s output files from Relate (.mut files) were combined and then separated by TE and SNP. TEs and SNPs that were flipped due to ancestral and derived allele mismatches, or were not mapped on unique branches, were removed from further analysis.

To validate the inferred genealogical trees along the genome, we compared the expected SFS for chromosome 1 from the inferred trees with the observed SFS for SNPs and for TEs (Figure S4). We calculated the expected SFS from the inferred trees by first simulating mutations onto the Relate-inferred ARG using msprime (Baumdicker *et al*. 2022) using a mutation rate map where we specified a mutation rate of 3.3e-8 (Clark *et al*. 2005) for unmasked regions and a mutation rate of 0 for masked regions. We then used the `allele_frequency_spectrum function` in tskit with `site` mode and `span_normalise` set to false (Wong *et al*. 2024; Kelleher *et al*. 2016; Ralph *et al*. 2020). Before calculating the observed SFS, we corrected the sites that were flipped by Relate due to ancestral and derived allele misspecification in the original input VCF, which included just 5694 SNPs and 439 TEs. We also dropped sites that did not map onto unique branches, which included a total of 30544 sites. Then, we calculated the SFS for SNPs and for TEs using the `sfs` function in scikit-allel (Miles *et al*. 2024). Finally, we length-normalized the SFS using the length of chromosome 1 minus the sum of the masked regions in base pairs. In addition, we calculated and plotted a normalized SFS to better compare the overall shape between the ARG and our data. The script to calculate and plot the SFS is available on GitHub (https://myersgroup.github.io/relate/parallelise.html,https://github.com/reginaanne/transposable_element_ages_selection).

We also compared the age (*T*) of each long terminal repeat (LTR) TE estimated from divergence (*K*) of its 5’ LTR and 3’ LTR sequences (SanMiguel et al. 1998), using the Kimura two-parameter method and the same mutation rate (*µ*) of 3.3 1× 0^−8^ substitutions per synonymous site per year, with the formula 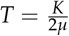 (Figure S4; Zhao et al. 2017).

### Φ_*SFS*_, ***a new statistic***

To account for the non-constant mutation rate of TEs, Horvath *et al*. (2022) developed an age-adjusted SFS by calculating the frequency difference (Δ_*f*_ ) between TEs and putatively neutral SNPs. However, interpreting the shapes of different TE groups is challenging (Figure S5). To improve this analysis, we developed a new statistic Φ_*SFS*_ to compare selection among different TE groups. As with the Δ_*f*_ approach, TEs were first binned into deciles based on their estimated age. Neutral SNPs were then downsampled to match both the number and age distribution of TEs within each age bin. Both TEs and SNPs were subsequently divided into 25 bins according to their allele frequency. Under neutrality, TEs are expected to accumulate at rates comparable to those of neutral SNPs with a matching age distribution, resulting in a frequency distribution where the proportional difference between the two is close to zero. However, selection on TEs will lead to a distorted SFS. In particular, selection against TEs is expected to lead to a left shift in the SFS, resulting in a higher density in lower allele frequency bins relative to SNPs. We quantified the overall SFS deviation of TEs by summing the proportional differences between TEs and SNPs across allele frequency bins using Equation 2, where higher Φ_*SFS*_ values indicate stronger deviations (and by implication, stronger selection).

### Simulation validation of Φ_SFS_

To test our Φ_*SFS*_ statistic, we performed simulations in SLiM

3.3.2 (Haller and Messer 2019a). We simulated a 10Kb region with a mutation rate of 10^−5^ and a recombination rate of 10^3^ in a constant-sized population of 1,000 diploids. Mutations were assumed to be either neutral or deleterious. We varied a number of parameters, including the proportion *p* ∈ (0.5, 0.6, 0.7, 0.8, 0.9, 1) of mutations that were selected, the dominance *h* ∈ (0.01, 0.25, 0.5) of selected mutations, and the selection coefficient *s* ∈ (10^−^3, 10^−^2, 10^−^1) of new mutations (Figure S6). We ran 100 iterations of each parameter combination for 10,000 generations, sampling 26 haploid genomes in the final generation and recording the frequency of all neutral and selected mutations separately.

### Burden of Rare Variants

We obtained gene expression data from the 26 NAM lines (Hufford et al. 2021) for each of ten tissue types: leaf mid, leaf base, leaf tip, shoot, root, tassel, anther, embryo, endosperm, and ear. Only genes with expression and annotations across all 26 NAM lines (Pan-genes) were retained for analysis (Hufford et al. 2021; Zeng et al. 2023). To investigate the effect of rare TEs on gene expression, we applied the rare variant burden test (Zhao *et al*. 2016) separately for each of the ten tissue types across the 26 NAM lines. For each gene, individuals were ranked by expression level and grouped into expression rank bins from low to high. The number of singleton TEs inserted in or near each gene was then summed across all genes within each expression rank bin. Under the null hypothesis, gene expression levels are independent of the count of singleton TEs. However, if singleton TEs contribute to extreme gene expression levels (either lowest or highest), a quadratic relationship is expected between the count of singleton TEs and expression rank. For all rare variant burden tests, we compared TEs located within genes to those located more than 5 kb away from genes.

### DNA methylation analysis

We downloaded whole-genome bisulfite sequencing (WGBS) data for leaf tissue from the 26 NAM lines from a previous study (Hufford *et al*. 2021) to investigate whether pre-existing DNA methylation levels at genomic sites influence the selection of transposable element (TE) insertions. CG methylation levels were measured in a 100 bp window surrounding each TE insertion site in NAM lines lacking the respective TEs (Figure S10A), serving as a proxy for the DNA methylation status prior to TE insertion. Based on methylation levels, TEs were classified into those inserted into highly methylated regions (methylation level ≥ 40%) and lowly methylated regions (methylation level ≤ 20%). To investigate whether the spreading of DNA methylation by TEs influences selection, we identified TEs that inserted into lowly methylated regions but were associated with increased DNA methylation following insertion (Supplementary Data; (Noshay *et al*. 2019)). Specifically, TEs were classified as spreading methylation if they inserted into regions with CG methylation ≤ 20% and, after insertion, either left or right flanking 100 bp region showed CG methylation ≥ 40%. In contrast, TEs inserted into regions with initial methylation ≤ 20% where flanking regions remained below this threshold after insertion were classified as not spreading methylation.

### Transcription Factor Impacts

To investigate whether TE insertions contribute to or disrupt transcription factor (TF) binding sites, we obtained MOA-seq data across 22 NAM lines from a previous study (Engelhorn *et al*. 2024). We first identified TF binding sites that overlap with TE insertion in each of the 22 NAM lines using BEDTools (Quinlan and Hall 2010). A TE was classified as overlapping if at least one NAM line was annotated as having a TF binding site that overlapped (1bp) with a TE insertion; otherwise, the TE was considered non-overlapping (Figure S11). The resulting TF binding site data were then merged across all lines to generate a polymorphic TF binding site matrix, representing the presence or absence of TF binding at each TE insertion across the NAM population.

Similarly, we extracted TE polymorphism data across the same 22 NAM lines to assess the presence or absence of each TE insertion within the population. By combining the polymorphic TF binding sites with TE presence/absence across the 22 NAM lines, we classified TEs into distinct categories based on their association with TF binding patterns (Figure S11). Negative: NAM lines with the TE insertion lack a TF binding site, while lines without the TE consistently have the binding site. Partial Negative: lines with the TE insertion lack the TF binding site, but among lines without the TE the binding site may or may not be present. Positive: TE-containing lines consistently have the TF binding site, while lines lacking the TE do not. Partial Positive: lines with the TE insertion have the TF binding site, but lines without the TE may or may not have it.

### TE expression analysis

To investigate how TE expression influences their selection, we obtained RNA sequencing data from 10 B73 maize tissue types from a previous study (Hufford et al. 2021). FASTQ files were aligned to the B73v5 reference genome, and TE expression was analyzed following the method described by Anderson *et al*. (2018). We modified our B73 TE annotation to meet HTSeq requirements and masked exon regions in the TE file using BEDTools (Quinlan and Hall 2010). Gene annotations for B73v5 were then integrated with the modified TE annotation file, and mapped reads were assigned to genomic features using HTSeq. The SAM output from HTSeq was analyzed combining both uniquely- and multi-mapped reads to generate per-family TE counts, and reads per million (RPM) values were calculated by normalizing to the total number of gene-assigned and TE family-assigned reads in each library (Anderson et al. 2018). For each tissue, genes were divided into four quartiles based on RPM. For each tissue, Φ_*SFS*_ values were compared between TEs in the first and fourth gene expression quartiles.

### Positive and balancing selection analysis

We used Selscan (Szpiech and Hernandez 2014) to identify potential TEs under positive selection across the genome. We first calculated the integrated haplotype score (iHS; (Voight et al. 2006)) for each variant in our VCF. We then used GenWin (Beissinger 2014) to partition the scores for TEs and SNPs into windows based on a spline fit to the iHS data. We identified outlier windows in the top 1 and 5% of iHS values by averaging across variants in each window. We compared the number of outlier windows in which the TE was the variant with highest iHS value to the number observed in randomly selected windows, performing the random sampling 100 times. Similarly, we ran BetaScan (Siewert and Voight 2017) to detect potential TEs under balancing selection by analyzing their *β* scores. Since negative *β* values are not biologically meaningful, they were removed from subsequent analyses. As before, we made windows with GenWin and compared the number of windows in which a TE had the highest *β* to those in randomly selected windows, performing the random sampling 100 times.

## Results

To study the dynamics of transposable element (TE) polymorphism in maize, we took advantage of high-quality assemblies of 26 maize genomes from the Nested Association Mapping (NAM) panel (Hufford et al. 2021; McMullen et al. 2009). Munasinghe *et al*. (2023a) combined TE annotations and structural variation (SV) data derived from alignments between each of the NAM lines and the B73 reference genome to identify TEs exhibiting presence-absence variation (“TE=SV” data; Figure S1). We linked the pairwise “TE=SV” across lines based on their position and size to obtain the presence/absence of each TE in all 26 genomes (Figure S2, see Methods). We also called single nucleotide polymorphisms (SNPs) between each of the NAM lines and the B73 reference genome based on these alignments and merged these pairwise comparisons with B73 to obtain a polymorphic SNP dataset. Our final data set included 140,250 TEs and ≈20M SNPs.

We divided the TEs into two groups based on their method of identification: structural TEs, identified by characteristic struc-tural features; and homology-based TEs, identified through sequence similarity to known elements. We further divided TEs into RNA (Class I) and DNA (Class II) based on their different transposition mechanisms. Among these 140,250 TEs, we observe 20,615 DNA-structural, 52,390 RNA-structural, 24,910 RNA-homology, and 42,345 DNA-homology TEs (Figure 1A). We annotated polymorphic SNPs using SnpEff (Cingolani *et al*. 2012), classifying them into four categories — High, Moderate, Modifier, and Low — based on predicted functional impact. Among the 20M SNPs, 394,949 (2%) are Low SNPs, primarily consisting of synonymous and intronic variants (Figure S3A).

**Figure 1.**
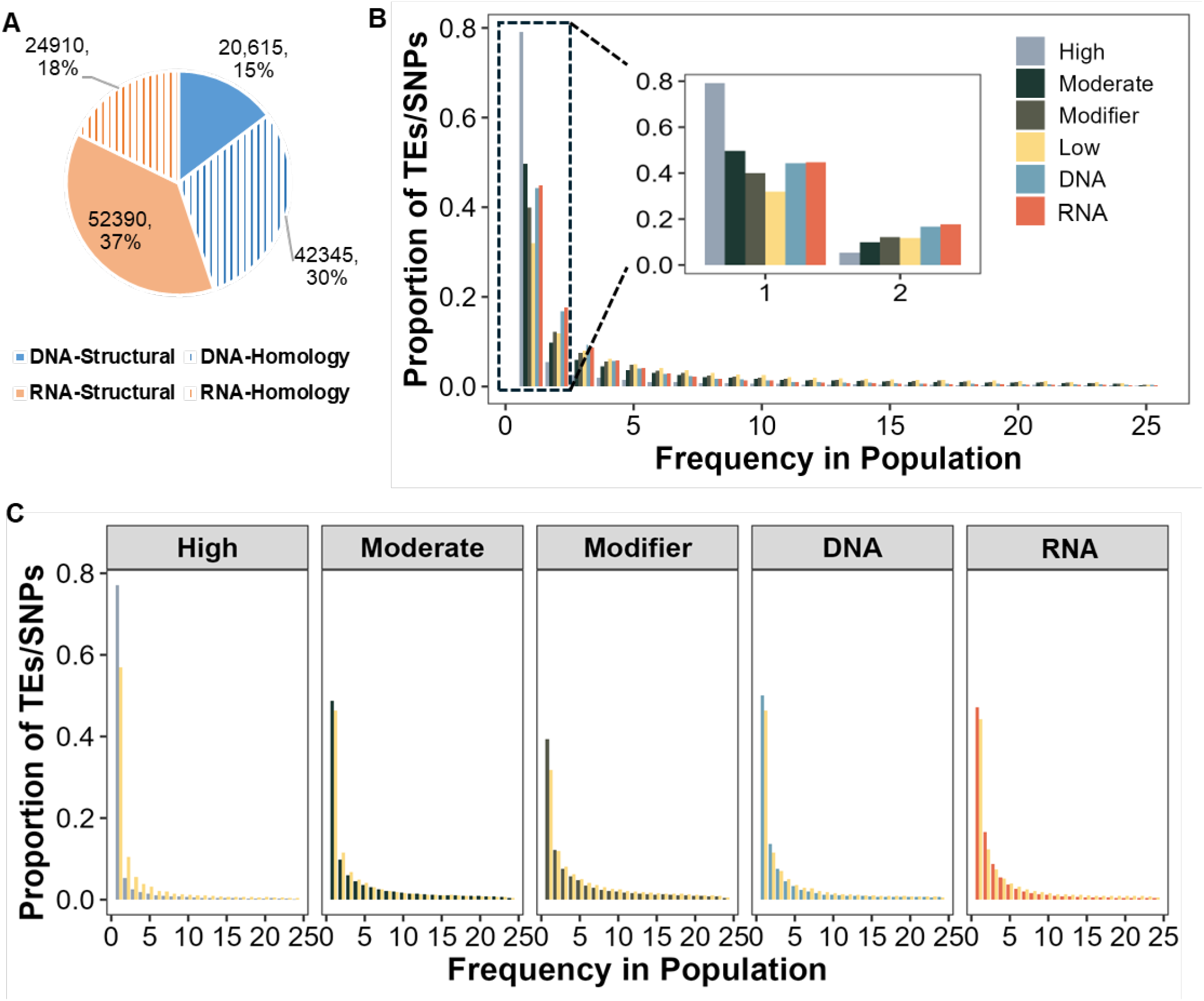
The importance of incorporating TE age in detecting selection using the site frequency spectrum (SFS). (A) Number and percentage of DNA and RNA TEs identified using structure- and homology-based methods. (B) Comparison of SFS distributions for TE groups and SNP categories identifiedd by SnpEff. (C) Comparison of SFS distributions for different SNP and TE groups with their corresponding age-adjusted Low SNP.

The site frequency spectrum (SFS) is widely used to detect signatures of selection acting on both SNPs (Casillas *et al*. 2007; Beissinger *et al*. 2016) and TEs (Horvath *et al*. 2024; Jiang *et al*. 2024; Kofler *et al*. 2012; Neafsey *et al*. 2004). In particular, variants affected by both negative and stabilizing selection typically show a skew towards rarer alleles than comparable neutral variants (Nielsen 2005; Charlesworth 2013). Consistent with these predictions, “High” SNPs (e.g., those annotated as disrupting gene function) exhibit the most pronounced left-skew, whereas “Low” SNPs showed the least (Figure 1B). We therefore use “Low” SNPs as the putatively neutral set for further comparisons. The SFS of both structural DNA- and RNA-TEs were more left-skewed than “Low” SNPs but less so than “High” SNPs (Figure 1B). In contrast, TEs identified purely through homology exhibited right-skewed distributions relative to Low SNPs (Figure S3B and S3C), perhaps due to difficulties identifying rare TEs using homology alone. We therefore excluded homology-based TEs from further analyses.

Because TE insertion dynamics are not uniform over time (Naito *et al*. 2006; Ungerer *et al*. 2009), formal comparison of TEs to SNPs must account for the age of each variant (Horvath *et al*. 2022). We estimate the age of each TE and SNP using the genealogy-based method RELATE (Speidel *et al*. 2019). We assign each TE or SNP an age based on the midpoint of the branch on which it is found. Comparison of the observed and expected SFS suggests branch lengths of the genealogies are somewhat overestimated (leading to a higher count of polymorphisms; Figure S4A-B). The relatively consistent shape of the SFS suggests that relative branch lengths (and thus relative ages of the variants) for much of the SFS are reflective of our data. These comparisons do highlight, however, that our genealogies overestimate the expected number of high-frequency derived variants, likely due to mis-polarization of variants in regions where we are lacking an outgroup alignment. Age estimates of LTR TEs derived from our genealogical approach showed only a weak correlation with ages estimated from pairwise differences between intra-element long terminal repeats (Figure S4C-D), likely reflecting potential biases in our approach as well as inaccuracies in estimates from substitution rates (see Discussion). Because age estimates cannot be directly estimated from DNA element sequences, phylogenetic branch-length approaches, not dissimilar from our ARG-based approach, are the only method available, and have been used in the past to estimate their ages (Stitzer *et al*. 2021).

### The Age-adjusted Site Frequency Spectrum

Comparison of both RNA and DNA TEs to a set of Low SNPs matched by age (see Methods) reveals less of a left-skew for TEs (Figure 1C), highlighting the importance of incorporating age into such comparisons. Horvath *et al*. (2022) introduced the age-adjusted SFS to account for TE age when evaluating evidence of selection. In this approach, TEs and SNPs are each binned into deciles by age, then neutral SNPs are downsampled to match the count and age distribution of each TE bin. The difference in mean allele frequency between TEs and SNPs, termed Δ_*f*_, is then plotted across age bin. TEs under negative selection show a characteristic pattern of lower frequencies — and thus a negative Δ_*f*_ — most evident in the oldest age bins. We first compared the Δ_*f*_ distributions across age bins for different SNP and TE groups, observing that High SNPs exhibited the most negative Δ_*f*_ (Figure S5). While the Δ_*f*_ profiles of both RNA and DNA TEs are suggestive of negative selection, the shapes of these distributions are difficult to interpret (Figure S5), likely due to variation in both the proportion of selected TEs and their fitness consequences.

To address difficulties in interpreting Δ_*f*_, we developed a new statistic, Φ_*SFS*_ that is both more powerful and easier to interpret, allowing effective comparison of selection across subsets of TEs in the genome. Following the same binning strategy used in Δ_*f*_ calculations, we grouped TEs into ten equal-sized age bins, downsampling Low SNPs to match the count and age distribution of each bin. We then pooled variants across all ages and compared the SFS of TEs and SNPs, calculating Φ_*SFS*_ as the sum of the positive differences in density between TEs and SNPs (Eq. 2; Supplementary Methods).

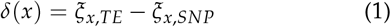

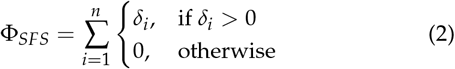

We first tested our Φ_*SFS*_ statistic with simulated mutations generated in SLiM (Haller and Messer 2019b), varying the dominance (*h*), selection coefficient (*s*), and proportion (*P*) of mutations under selection. As expected, Φ_*SFS*_ was most sensitive for mutations with additive effects on fitness, though still performed well even for the recessive case. With strong selectio (2*Ns* = 200), polymorphisms are kept at such low frequency that a large proportion of the variants observed in a sample size similar to our data are neutral, and Φ_*SFS*_ loses some of its power (Figure S6). Under all scenarios, however, Φ_*SFS*_ decreases as the proportion of SNPs under selection decreases (Figure S6), reaching ≈ 0 when *P* = 0 and all mutations are neutral.

We then used Φ_*SFS*_ to compare genome-wide TE and SNP data, using 100 bootstrap samples of age-matched samples of Low SNPs to generate confidence intervals. Consistent with the Δ_*f*_ results, we observed that High SNPs had the highest Φ_*SFS*_, while DNA and RNA elements exhibited intermediate values (Figure 2A).

**Figure 2.**
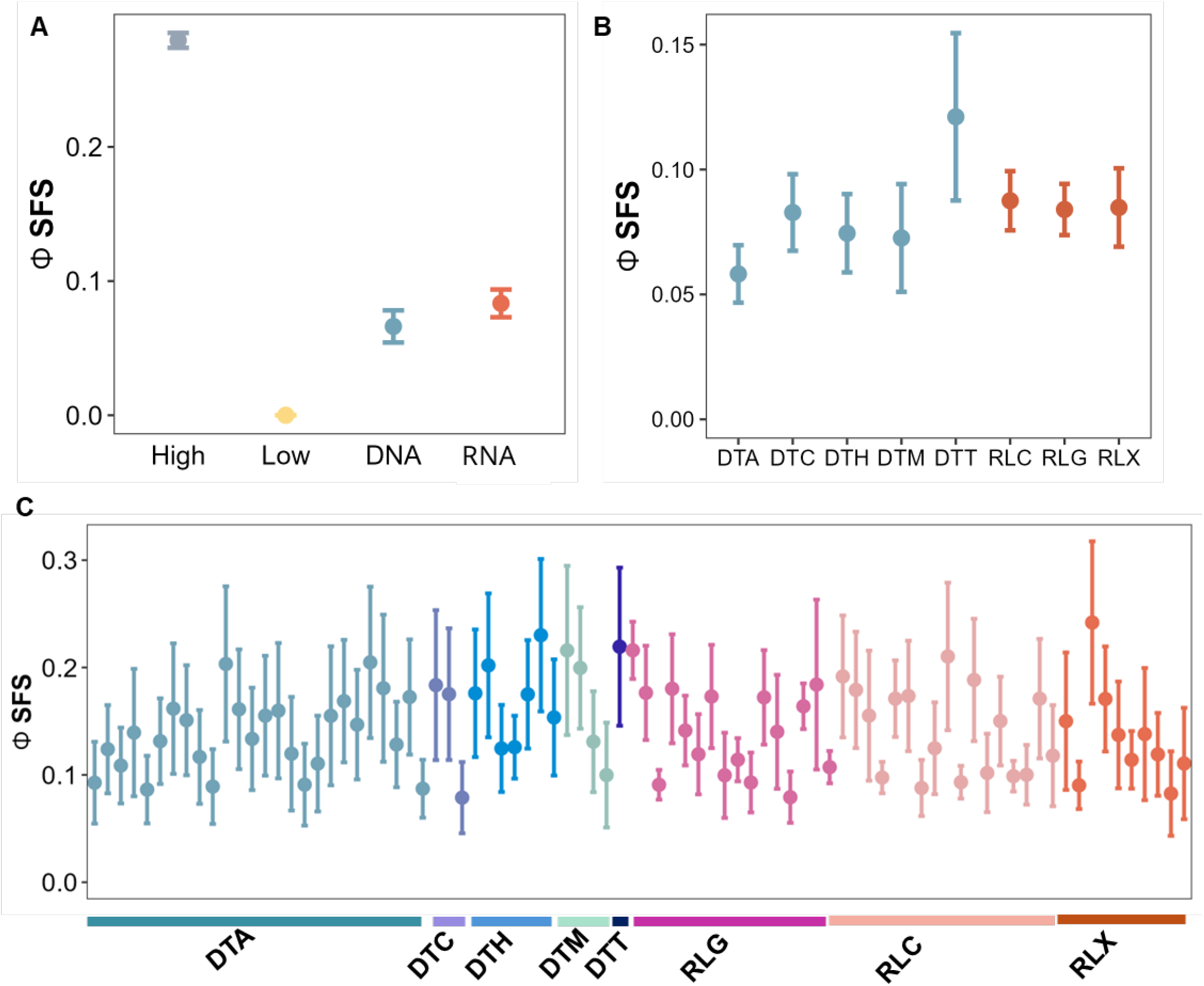
Φ_SFS_ captures differences in selection among SNPs and TEs at multiple classification levels. (A) Selection on different SNP (high and low effects on gene) and TE (DNA and RNA) groupw. (B) Selection across TE superfamilies. (C) Selection among TE families with more than 100 insertions.

In addition to the broad DNA and RNA distinction reflecting their mode of transposition, TEs can be further classified into superfamilies and families based on sequence similarity (Wicker et al. 2007). Different families and superfamilies in the maize genome show diverse histories, behavior, and relationships with the host genome (Stitzer et al. 2021), suggesting that not all should be equally affected by selection. We divided RNA and DNA TEs into eight superfamilies, with the majority belonging to RLG, RLC, and DTA (Table S1). We observed similar Φ_*SFS*_ values among RNA superfamilies, but greater variation among DNA elements, with DTA showing the weakest evidence of selection, the CACTA (DTC) superfamily showing a Φ_*SFS*_comparable to RNA TEs and DTT showing the highest Φ_*SFS*_(Figure 2B). Most TE families had higher values of Φ_*SFS*_compared to that of their corresponding superfamilies, with some families such as CRM2 (RLG), Tourist (DTM) with high Φ_*SFS*_ comparable to loss-of-function SNPs (Figure 2C). The difference in Φ_*SFS*_ between families and their corresponding superfamilies may reflect a loss of power at the family due to averaging of across multiple families within each superfamily.

### Location Matters

Perhaps the clearest potential detrimental effect of TEs is their ability to disrupt normal gene function. We observed that Φ_*SFS*_ decreases with increasing distance between TEs and genes for both DNA and RNA elements (Figure 3A). As we saw previously, RNA TEs continue to exhibit stronger signatures of selection compared to DNA TEs across all genomic locations. To assess whether the negative relationship between Φ_*SFS*_ and TE distance from genes is robust to differences in linked selection between genes and intergenic regions (Beissinger et al. 2016), we compared the Φ_*SFS*_ of intergenic SNPs far from genes (identified by SNPeff as “modifier” SNPs and representing 96% of all SNPs) in different genomic locations. In contrast to TEs, we found that the Φ_*SFS*_ of intergenic SNPs increases with increasing distance between SNPs and genes (Figure S7). This pattern is inconsistent with a model of background selection (Beissinger et al. 2016), but suggests differential selection on TEs than on intergenic SNPs in our data.

**Figure 3.**
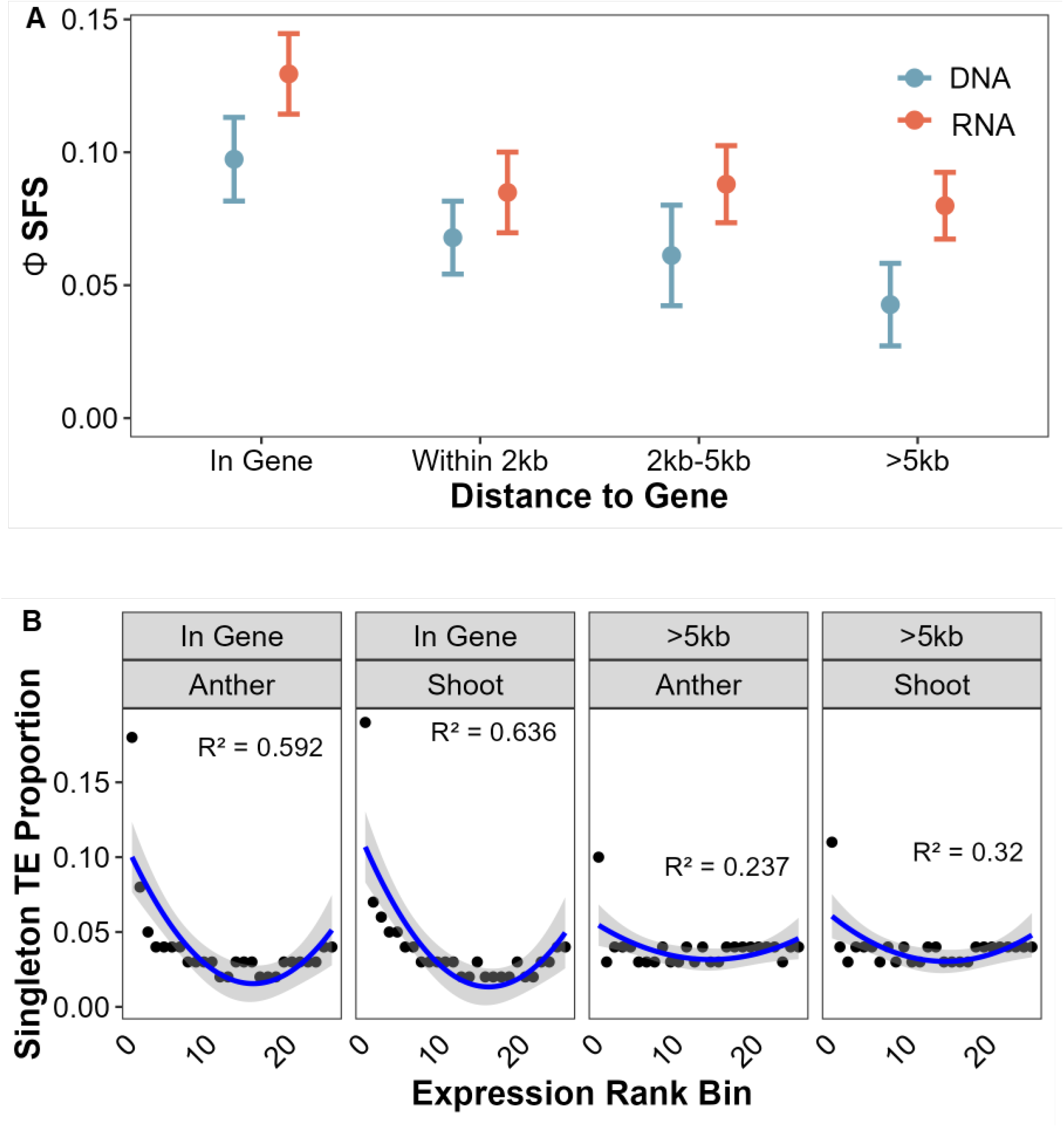
Location is the most important factor for TE selection. (A) selection on DNA and RNA TE across different genomic locations. (B) Relationship between TE count and gene expression rank bins for rare variants (singleton TEs) inserted within genes > 5kb from genes, shown for anther and shoot tissues.

If TE insertions in or near genes are subject to stronger selection due to their potential effects on nearby gene expression then they should be kept at low frequency, and genes exhibiting aberrant expression should be more likely to harbor rare TEs (Uzunovicét al. 2019; Kremling et al. 2018). To test this hypothesis, we analyzed gene expression data from 26 NAM lines across 10 tissues. Consistent with this prediction, we find that, across genes, the NAM line with the lowest expression had a notably higher count of singleton TEs, both within the gene model and more than 5kb away (Figure 3B and Figure S8). We also observed that singleton TEs were enriched in genes at the highest expression bin, suggesting selection is acting to reduce the frequency of TEs causing any large variation in expression.

### Epigenetic and Regulatory Impacts of TEs

Our results strongly support the idea that TEs near genes impact gene function. These observations could be due to the direct impacts of insertions into conserved coding or regulatory sequence (Noshay *et al*. 2021; Engelhorn *et al*. 2024), or the effects of epigenetic silencing of TEs after insertion (Lisch 2013; Huang and Lee 2024). Due to their deleterious impact, most TEs are regulated by the host through various mechanisms, among which DNA methylation plays a central role (Lisch 2009; Liu and Zhao 2023). Although methylation primarily serves to silence TEs, changing methylation levels induced directly by silencing of TE insertions or indirectly by the spreading of methylation into adjacent regions may have unintended side effects on gene expression. We first hypothesized that, while TE bodies are often highly methylated (Figure S9A), this may be less problematic if the TE inserted into a region that was already highly methylated. To test this hypothesis, we analyzed whole-genome bisulfite sequencing (WGBS) data for the 26 NAM lines (Hufford *et al*. 2021). As a proxy for the methylation status of the region prior to TE insertion, we calculated the average CG methylation level within a 100bp window surrounding TE insertion sites across NAM lines lacking the TE insertion (Figure S10A). Consistent with our hypothesis, we observed that, for TEs located in the same genomic regions, those inserted into lowly methylated regions exhibited higher Φ_*SFS*_ values compared to TEs inserted into highly methylated regions, a trend that was particularly clear for DNA TEs (Figure 4A). Moreover, for TEs inserted into the same methylation category, the Φ_*SFS*_ decreased as the distance between the TE and the nearest gene increased (Figure 4A), consistent with the patterns reported in Figure **??**A. We then hypothesized that TEs which induce methylation spreading into adjacent genes should experience stronger negative selection than those that do not. Contrary to our hypothesis, however, we saw no consistent difference in Φ_*SFS*_ between TEs differing in patterns of methylation spreading (FigureS10B).

**Figure 4.**
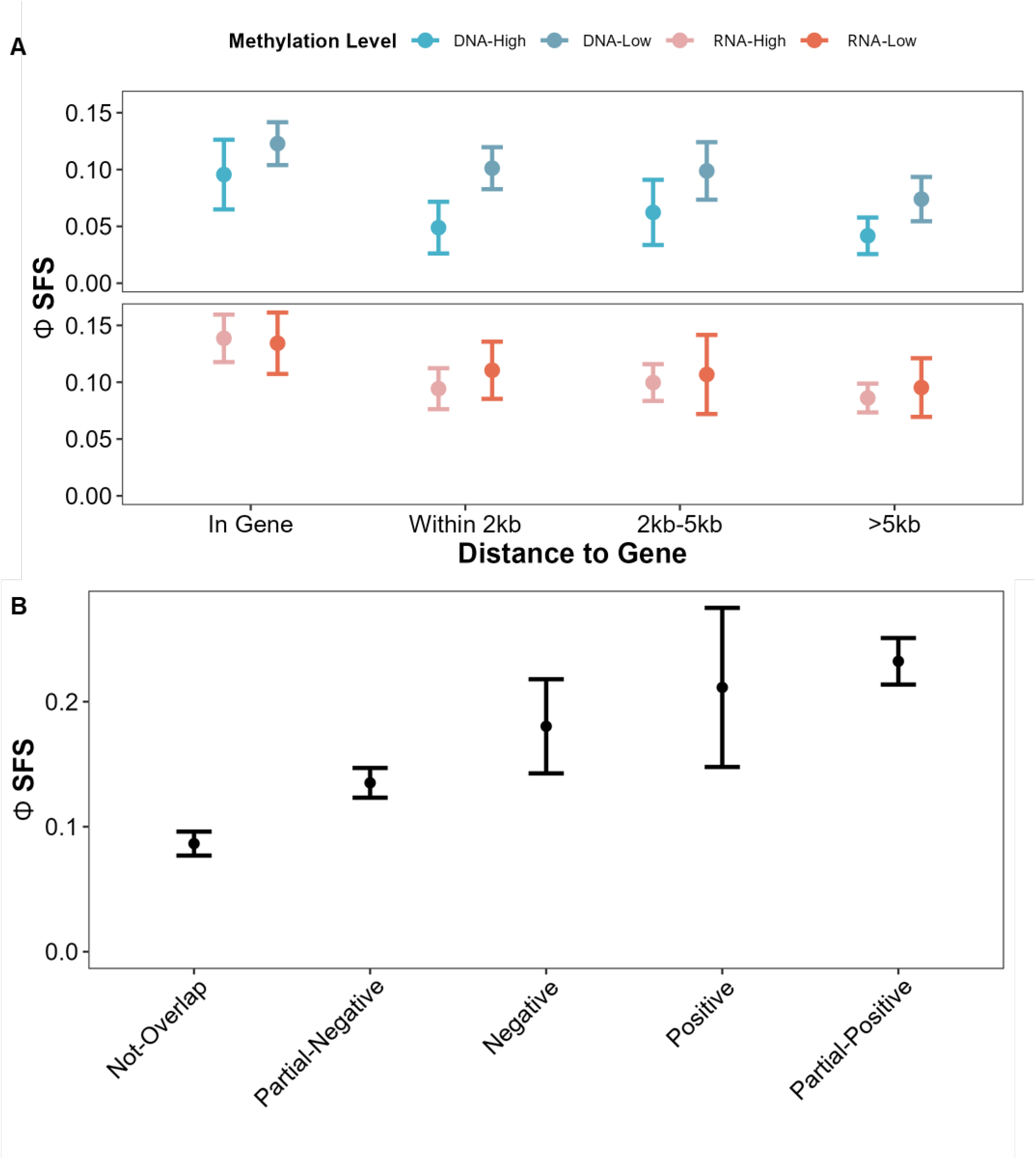
Selection on the epigenetic and regulatory changes induced by TEs. (A) selection on DNA and RNA TEs inserted into highly ( ≥ 0.4) or lowly ( ≤ 0.2) methylated regions across different genomic locations.(B) Selection on TEs categorized by their relationship with transcription factor (TF) binding sites, based on their association with a gain (positive), loss (negative), or no change (not-overlap) in TF binding sites.

Since our results indicate that TEs inserted into unmethylated regions are subject to stronger selection, we hypothesize that this may be due to TEs affecting regulatory elements located within these unmethylated regions. Although there are documented cases in which TEs contribute to the formation of novel transcription factor (TF) binding sites and also disrupting existing TF binding sites (Noshay *et al*. 2021; Studer *et al*. 2011), the frequency with which TE insertions either create or disrupt these regulatory elements on a genome-wide scale remains unclear. Recent studies characterizing TF binding sites in NAM lines have revealed frequent overlap between TEs and TF binding sites, including several examples where TEs disrupt binding sites and downstream gene regulation (Engelhorn *et al*. 2024). We would expect that TE insertions that create or disrupt TF binding sites are more likely to impact gene expression and thus be under stronger selection than those that do not. To test this hypothesis, we obtained presence/absence data of TF binding sites across 22 NAM lines (Engelhorn *et al*. 2024) and merged these with our TE polymorphism data (Figure S11). In total, we identified 15,855 TEs overlapping with TF binding sites and 57,150 TEs not overlapping with TF binding sites (Table S2). Among the 15,855 TEs, 376 exhibited a perfect negative association with TF binding sites, followed by more than 11,500 that showed a partial negative association (Figure S11). A much smaller number of TEs showed positive (117) or partial positive (2,554) associations with TF binding sites, respectively (Table S2), suggesting that TE insertions predominantly disrupt TF binding sites on a genome-wide scale. More importantly, we observed that TEs exhibiting both negative and positive relationships with TF binding sites are subject to stronger selection compared to those non-overlapping groups (Figure 4B). Notably, the magnitude of Φ_*SFS*_we observe for TEs that affect TF binding sites is higher even than the Φ_*SFS*_of TEs in genes, and approaches that of SNPs known to disrupt protein function (Figure 2).

### Features of Transposable Elements

We next examined how intrinsic characteristics of TEs contribute to the selective pressures they experience. First, we examined the impact of copy number, which varies significantly among different classes, superfamilies, and families (Stitzer et al. 2021) Although the relationship between TE copy number and host fitness remains unclear, several hypotheses merit further investigation. One hypothesis, supported by some empirical evidence (Huang and Lee 2024), is that TE insertions exhibit synergistic epistasis, such that each additional insertion has a larger impact than the last. This hypothesis predicts weaker selection on TE families with higher copy numbers, either because synergistic epistasis only allows weakly selected families to reach higher copy number or because more effective silencing from small RNAs reduces the insertional mutagenesis of high copy families. In contrast, higher copy number families may experience stronger selection because they lead to a greater risk of deleterious ectopic recombination (Kent *et al*. 2017), which could threaten genome integrity, or because they exact a higher metabolic cost to the host genome (Badge and Brookfield 1997).

Separating families by their copy number for both DNA and RNA TEs inserted in the B73 reference genome, we find that larger (>= 10 copy) families exhibit higher Φ_*SFS*_ values for RNA TEs (Figure 5A), but little difference among DNA TEs regardless of their family size. If stronger selection on large RNA families was due to their metabolic cost, we would expect to see that higher-copy families exhibit more transcripts (and thus greater cost to the host), but we see no meaningful correlation between copy number per family and expression (Figure S12).

If ectopic recombination helps explain selection against high-copy RNA families, we might also expect to see an impact on TE length. Longer TEs are more likely to participate in illegitimate recombination events, and thus subject to stronger negative selection (Song and Boissinot 2007). To test this hypothesis, we stratified DNA and RNA TEs into four length categories

(Figure S13). While we did not observe the predicted negative relationship between Φ_*SFS*_ and TE length for either DNA or RNA TEs, even after controlling for the distance between gene and TE (FigureS13), we do note that the length distribution of TEs shifts towards longer copies farther away from genes (Figure 5B), and note lower Φ_*SFS*_values for small RNA elements in low recombination regions of the genome (Figure S14).

**Figure 5.**
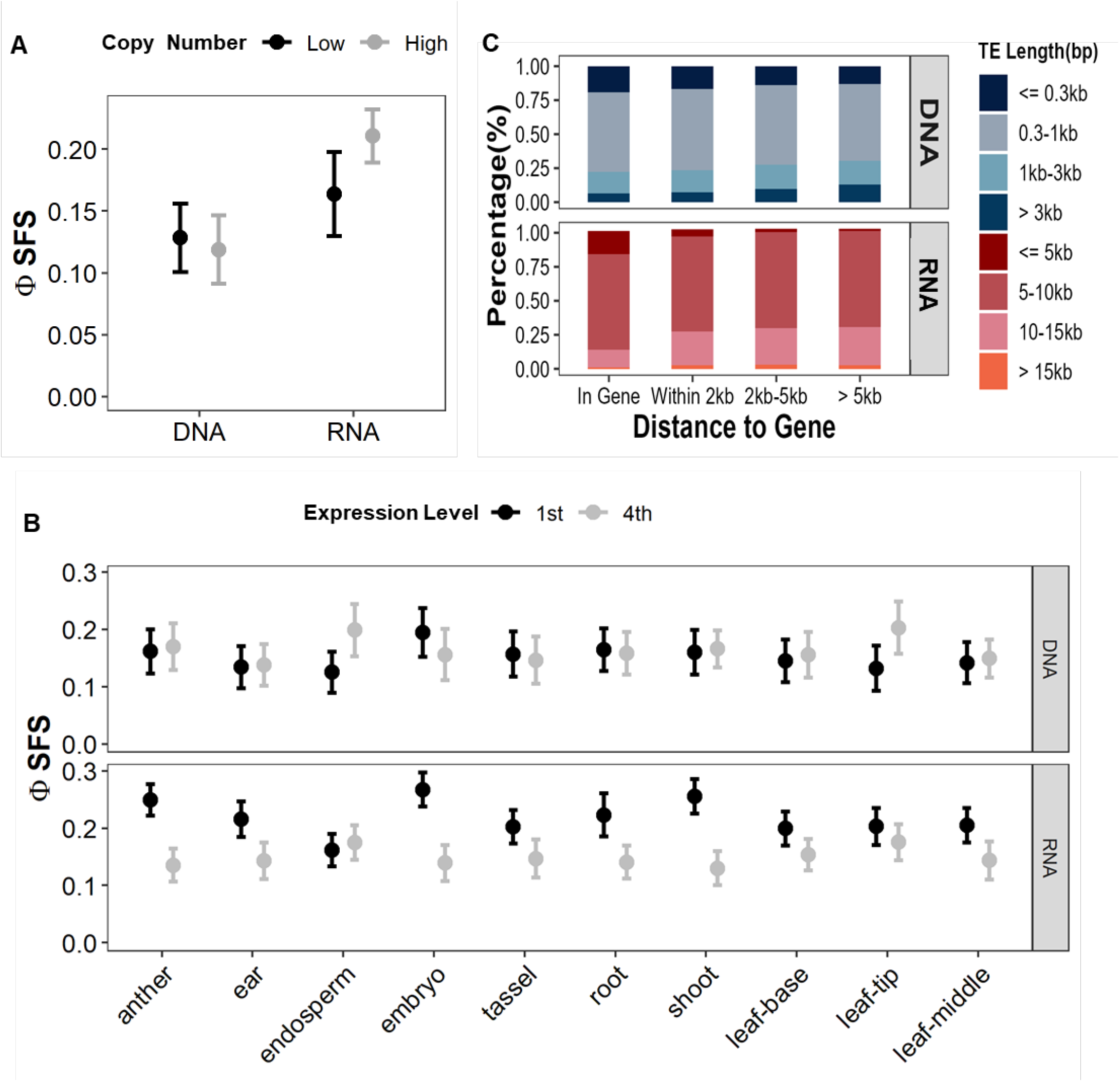
Effects of intrinsic TE features on selection. (A) Selection on DNA and RNA TE families with low (< 10 copies) and high ( ≥ 10 copies) copy numbers.(B) Selection on TEs grouped by expression level across 10 tissues in B73, categorized into quartiles (1^st^ = lowest expression, 4th = highest expression).(C) Length distribution of DNA and RNA TEs across different genomic locations.

Transcription of TEs is essential for the production of proteins required for their transposition, and we hypothesized that more highly expressed TEs may thus be subject to stronger selection due to their increased mutagenic potential . However, small RNAs that initiate TE silencing are also derived from TE transcripts. Thus, higher TE expression may lead to increased small RNA production and more effective TE silencing, potentially resulting in weaker selection. To test these two different hypotheses, we analyzed RNA sequencing data from 10 tissue types of B73 and mapped them using an approach that ameliorates much of the multi-mapping challenges associated with repetitive DNA (Anderson et al. 2018). Consistent with the prediction of our second hypothesis, RNA TEs with lower expression exhibited higher Φ_*SFS*_ compared to TEs with higher expression across most of the ten tissues evaluated (Figure 5B). However, the selection are similar for DNA TEs regardless of expression across most tissue types.

### Positive and Balancing Selection

Although most TEs are likely neutral or deleterious because of their large impacts on phenotype, a subset may confer adaptive advantages and be subject to positive or balancing selection. To explore this possibility, we first identified windows with high iHS values, a statistic indicative of recent positive selection (see Methods). While we found a number of putatively selected windows in which the variant with the highest iHS value was a TE, this did not occur at a frequency higher than found in random windows (Figure S15A-C), providing little evidence that TEs are enriched for targets of positive selection. We performed a similar genome-wide scan to detect balancing selection using BetaScan (Siewert and Voight 2017) and likewise found putatively selected TEs at a frequency no higher than expected from random windows (Figure S15B-D).

## Discussion

In this study, we present the first population genetic analysis of transposable element (TE) polymorphisms in maize using highquality long-read genome assemblies. Leveraging the genome assemblies of the 26 parents of the Nested Association Mapping (NAM) population, we investigated TE insertion dynamics and the evolutionary forces shaping their distribution. We developed a novel statistic, Φ_*SFS*_, which incorporates TE and SNP age and offers improved interpretability over the previously used Δ_*f*_ (Horvath et al. 2022) (Figure 2A, Figure S5). Using Φ_*SFS*_, we detected variation in selection across TE classifications (Figure 2) and identified key factors influencing these patterns. Notably, TE insertion location emerged as a major determinant: selection strength decreased with increasing distance from genes (Figure 3A), perhaps due to effects on gene expression (Figure 3B). Consistent with this, regulatory changes associated with TE insertions contributed to variable selection on TEs (Figure 4). Finally, we observed that some intrinsic TE features, such as copy number and family-level expression, also influenced selection (Figure 5).

### The new statistic Φ_SFS_

Because TEs accumulate more sporadically than single SNP mutations, differences in allele frequencies between the two are difficult to interpret without knowing the age of individual variants. While an age-adjusted SFS method was previously developed to account for this issue (Horvath et al. 2022), it remains difficult to interpret when comparing selection across different TE groups (Jiang *et al*. 2024; Horvath *et al*. 2024). To address this limitation, we developed a new statistic, Φ_*SFS*_, designed for more intuitive comparisons among TE groups. We first validated Φ_*SFS*_ using simulated data and found it reliably distinguished differences in the proportion (*p*) of selected mutations across scenarios (Figure S6). However, under strong selection (e.g., *N*_*e*_*s* = 200) with additive mutations (*h* = 0.5), Φ_*SFS*_ showed reduced sensitivity. This is expected, as strongly deleterious mutations are rapidly purged, leaving only weakly deleterious or neutral variants at high enough frequency to be observed in a relatively small sample. Applying Φ_*SFS*_ to real data, we observed improved resolution in detecting selection differences among TE groups compared to Δ_*f*_ (Figures 2 and S5). While Φ_*SFS*_ does not directly estimate the proportion of TEs under selection, its relative values can be meaningfully compared across groups, with higher values indicating stronger selection (Figure S6).

### Genealogy and neutral SNP caveats

Here we use RELATE Speidel *et al*. (2019) — a genealogy-based method to estimate the age of both SNPs and TEs. Visualization of the SFS predicted from the estimated trees finds that RELATE predicts more mutations than seen in our empirical data (Figure S4A), likely due to mis-specification of the mutational target size. When comparing the shape of the density of the SFS, we found that the overall pattern is similar for low to intermeidate frequency derived SNPs, but trees estimated from RELATE still overestimate the density of high-frequency derived variants (Figure S4B). Because we are concerned with the relative age of SNPs and TEs, and this bias is strong only for rare, highfrequency derived alleles, we argue it should not have a major impact on our inference of selection against TEs.

We also compared the age of LTR elements estimated using RELATE to that using the sequence identify between the two LTRs of each TE. These measures showed little correlation (Figure S4C-D). There are several reasons for this, including our use of branch midpoints to estimate age as well as challenges in estimating age from LTRs — including mutations that arise during insertion (Preston 1996), the poor resolution to estimate young ages with few mutations, and potential annotation errors in the long terminal repeats themselves.

Another caveat of our analysis is the set of SNPs used as a neutral null. Identifying neutral SNPs is essential for using the site frequency spectrum (SFS) to detect selection. Previous studies applying the SFS to TEs often use synonymous SNPs as proxies for neutrality (Horvath *et al*. 2024; Jiang *et al*. 2024). Tools like SnpEff (Cingolani *et al*. 2012) classify SNPs based on their predicted impact on gene function — categorized as High, Moderate, Low, or Modifier. We define “neutral” SNPs as those with Low predicted effects, which were primarily synonymous variants. However, in maize, even synonymous SNPs may be linked to selected sites within genes, and thus not reflect neutral frequency distributions (Beissinger *et al*. 2016). We refrained from using intergenic SNPs as an alternative, as our filtering for sites present in all samples likely biases intergenic sites toward conserved, potentially functional regions also under selection (Casillas *et al*. 2007).

### Location, location, location

The genomic location of TEs plays a critical role in their silencing and evolutionary dynamics (Sigman and Slotkin 2016). TE insertions can alter gene function or regulation, particularly when located within or near genes (Bourque et al. 2018; Baduel *et al*. 2021). By characterizing both RNA and DNA TEs based on their proximity to genes, we found that TEs located within or near genes experience the strongest selection (Figure 3A), consistent with previous findings (Hollister and Gaut 2009). Because decreasing Φ_*SFS*_farther away from genes could be the result of linked selection rather than direct selection on TEs, we compared these results to the frequency spectrum of intergenic (modifier) SNPs. Rather than mirror the TE data as would be expected if our results were driven by linked selection, Φ_*SFS*_actually increases farther away from genes (Figure S7), perhaps reflecting biases in our stringent SNP filtering that may enrich for SNPs in conserved non-coding regions. Therefore, our finding that selection on TEs decreases with increased TE-gene distance is unique to TEs and not simply a function of linked selection.

We next investigated why TEs near genes are subject to increased selection. It is widely believed that such TEs experience stronger selection due to their potential impact on gene expression. Consistent with this hypothesis and previous studies Kremling *et al*. (2018); Uzunovicé*t al*. (2019), our results show that rare TEs are enriched in genes with extreme expression levels (Figure 3B).

### Epigenetic and regulatory changes induced by TE

Although 85% of the maize genome consists of TEs, the vast majority are silenced by the host genome. Among the various mechanisms employed by the host to silence TEs, DNA methylation plays a major role.(Lisch 2009; Liu and Zhao 2023). However, TE silencing can also affect gene expression through the spreading of DNA methylation to adjacent regions (Lisch 2013; Liu and Zhao 2023). Previous research (Hollister and Gaut 2009; Horvath and Slotte 2017) showed that methylated TEs near genes are under stronger negative selection than unmethylated ones. In maize, however, it is more complicated, in that most TEs are methylated (Figure S9) and TE methylation could results either from insertion into regions already highly methylated or insertion into previously unmethylated regions followed by de novo methylation. Using whole-genome bisulfite sequencing (WGBS) and TE polymorphism data from 26 NAM populations, we showed that TEs inserted into low-CG methylated regions are under significantly stronger negative selection than those inserted into already highly methylated regions, across all genomic contexts (Figure 4A). Although it has been widely recognized that methylation spreading of TEs can affect adjacent gene expression (Huang and Lee 2024), we find no evidence that selection differs meaningfully between TEs that spread methylation to neighboring regions and those that do not (Figure S10B). This may be because only certain TE families exhibit methylation spreading (Choi and Purugganan 2018) and our analysis pooling different families could obscure signals of selection. Nonetheless, our results suggest that TE inserted into unmethylated regions are likely more harmful for the host and therefore under stronger than those that inserted into already highly methylated regions (Figure 4A).

Alternatively, TE insertion near genes may impact the regulatory features such as transcription factor (TF) binding sites (Engelhorn et al. 2024; Bubb et al. 2025). We found that TEs either disrupting or creating TF binding sites are under significantly stronger selection than TEs with no association with TF binding (Figure 4B). As most TF binding sites are unmethylated, this is consistent with out result that TE insertion unmethylated regions are under stronger selection.

### Intrinsic characters of TEs

In addition to TE-host interactions, intrinsic features of TEs—such as copy number, length, and expression—can influence their selective dynamics. We found that RNA TE families with high copy number and low expression are under stronger negative selection (Figure 5A-B). Several hypotheses have been proposed to explain selection on transposable element (TE) families with varying copy numbers. One suggests that TE may be deleterious to the host genome due to the fact that their repetitive increases the likelihood of ectopic recombination (Kent *et al*. 2017), with higher copy numbers incurring greater costs and thus experiencing stronger negative selection. Another posits that small RNA-mediated silencing increases with TE copy number, leading to stronger methylation and weaker selection due to reduced TE activity (Huang and Lee 2024; Betancourt et al. 2024). Our finding that larger RNA TE families are under stronger selection than those small families is consistent with the first hypothesis that more TE increase the the likelihood of ectopic recombination and therefore under stronger selection (Figure 5A). However, our finding that RNA TE families with higher expression are under weaker selection is consistent with our second hypothesis that higher TE expression might lead to more small RNA generation, therefore stronger silencing of TE and weaker selection on them. Figure5B

Another character of TE that could affect the likelihood of ectopic recombination is the length of TE. Ectopic recombination is positively associated with local recombination rate (Kent *et al*. 2017), and longer TEs are more likely to participate in ectopic recombination events (Petrov *et al*. 2011). Given the potential deleterious effects associated with such events, we predicted that longer TEs would be subject to stronger negative selection than shorter ones. While we did not observe consistent selection on TE length, we did find that TEs near genes were shorter than TEs farther away from genes (Figure 5C), consistent with the notion that longer elements are more prone to ectopic recombination – and thus more likely to be removed – in regions of higher recombination (Song and Boissinot 2007). Perhaps consistent with this interpretation, we see that shorter RNA TEs appear to be under relaxed selection in low recombination regions of the genome, but long RNA TEs are under higher selection at regions with both high and low recombination rate (Figure S14) . The effect of length on the selective consequences of TE insertions might partially explain our observation that RNA TEs, which are generally longer (Figure S13B-C) than DNA TEs, are under stronger selection (Figure 3A).

### The Mode of Selection on TEs in Maize

While individual examples of beneficial TE insertions have been identified in maize (Huang *et al*. 2018; Studer *et al*. 2011), we find little evidence of positive or balancing selection on TEs genomewide (Figure S15). Our Φ_*SFS*_statistic detects deviation from the SFS, but Φ_*SFS*_alone is not informative of the mode of selection. In most cases this appears to be selection against TEs, leading to a left-shifted SFS (Figure 1). This has been traditionally been interpreted as negative selection against unambiguously deleterious TEs (Horvath *et al*. 2024; Jiang *et al*. 2024). We argue instead this is likely due to stabilizing selection, where the strength of selection scales with the phenotypic effect of an allele. For example, we found that rare TEs inserted in genes are enriched in genes with extreme (both lowest and highest) expression (Figure 3B), consistent with previous results for TEs in *Capsella Grandiflora* (Uzunovicé*t al*. 2019) and rare SNPs in maize Kremling *et al*. (2018). Similarly, TEs that either disrupt or create TF binding sites are under stronger selection compared to those that do not overlap with TF binding sties (Figure 4B). Overall, this suggests a picture where TEs are sslected against because of their large impact on phenotype (and thus fitness), not because they are unambiguously deleterious in all environments. Indeed, the large effect of TEs on phenotype may be why they are often found underlying adaptations to dramatically new environments (Yang *et al*. 2013; Huang *et al*. 2018; Hof *et al*. 2016; Aminetzach *et al*. 2005).

## Acknowledgments

We would like to thank Michelle Stitzer for helpful discussion and Julia Engelhorn for sharing early access to transcription factor binding data. We would like to thank Felix Andrews for bioinformatics help switching the rows and columns of our data. We acknowledge funding from NSF (grant 1934384) and USDA Hatch project CA-D-PLS-2066-H 548. This research used the High-Performance Computing Core Facility (HPC@UCD) at the University of California, Davis.

---

**Figure S1.**
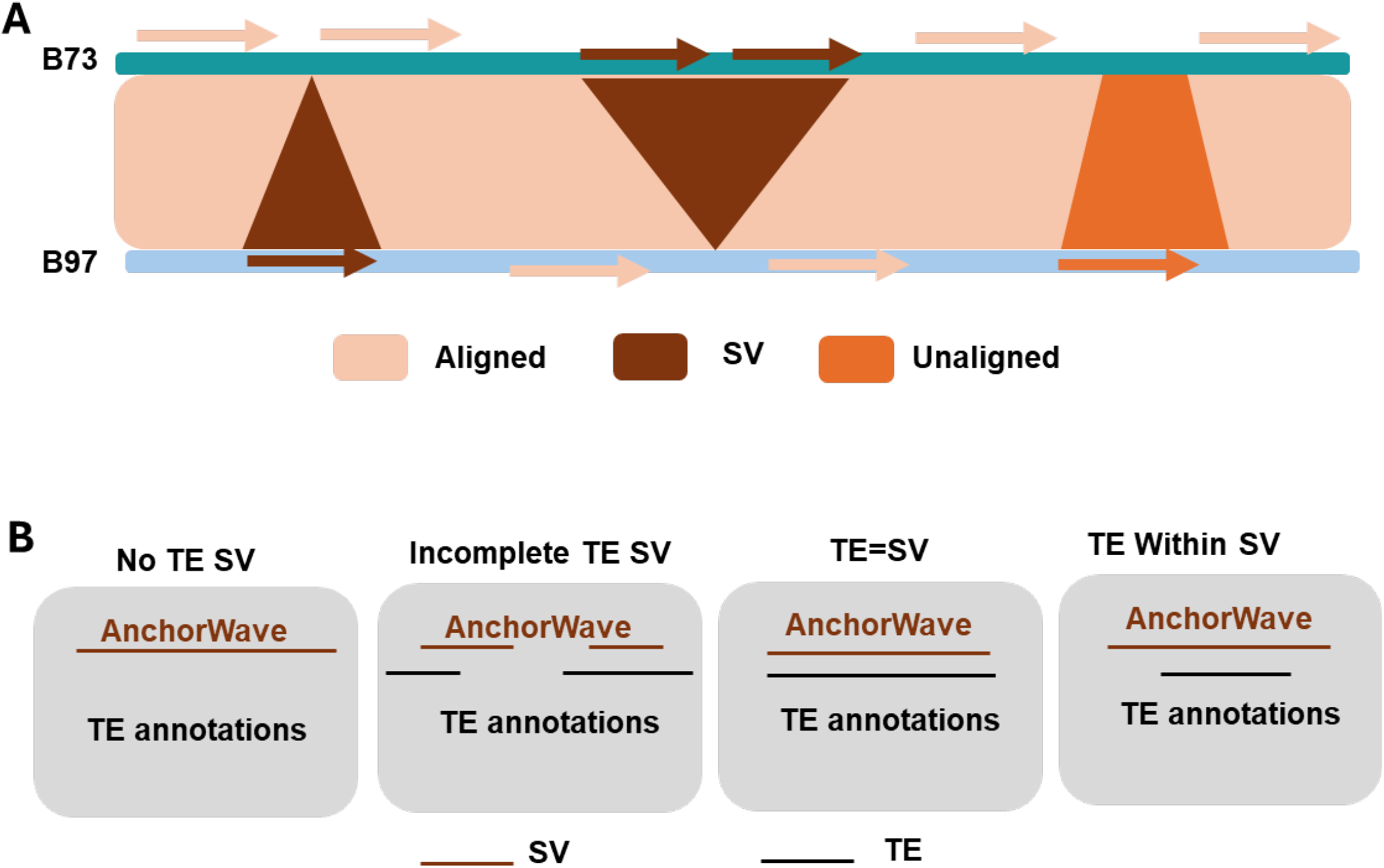
Identification of “TE = SV” Events. (A) A schematic illustrating the logic used to partition pairwise AnchorWave alignments between B73 and each NAM founder line into alignable regions, structural variants (SVs), or unalignable sequences.(B) A diagram showing how SVs were classified based on their overlap with transposable element (TE) content.

**Figure S2.**
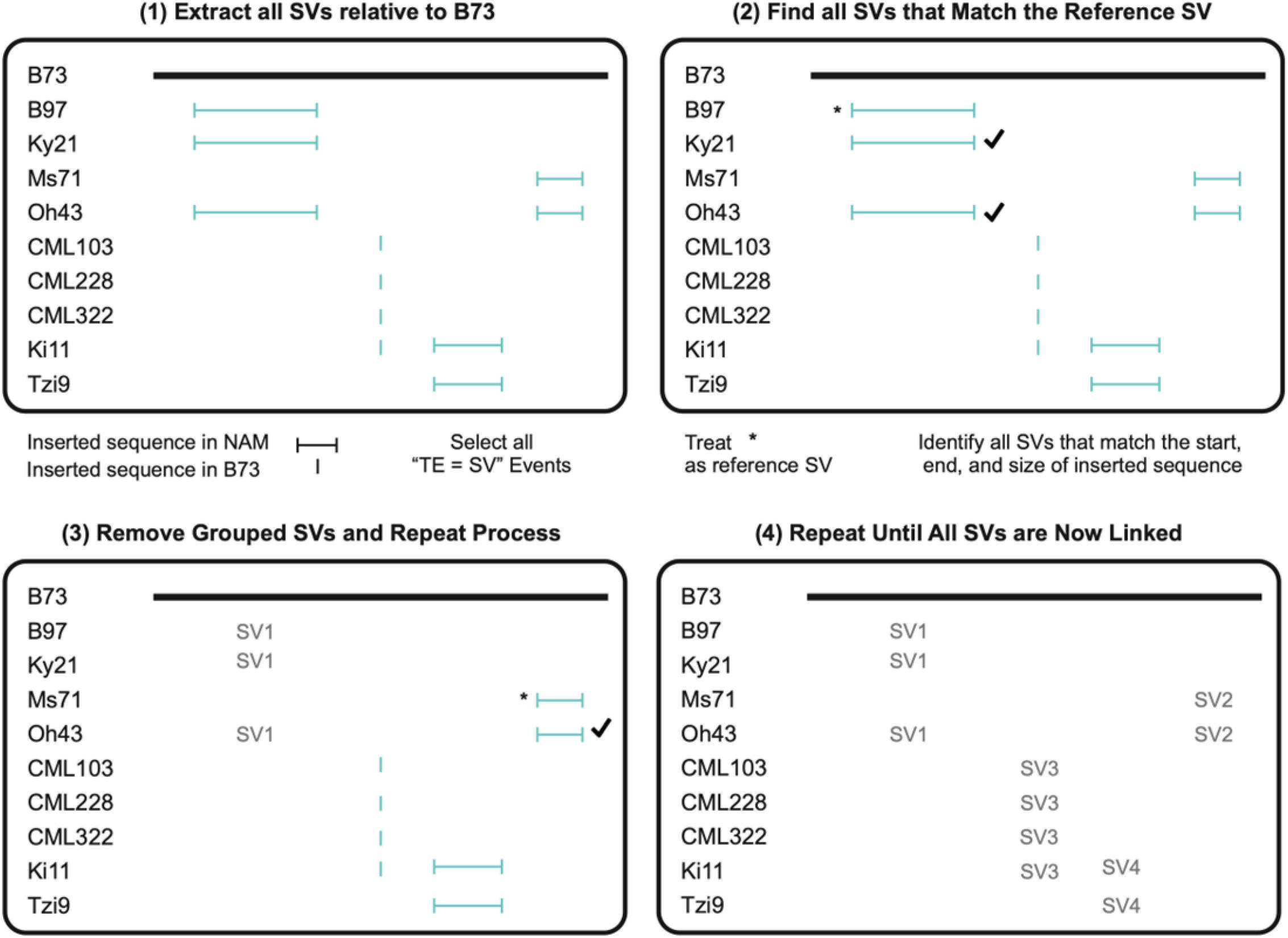
Linking “TE = SV” Events Across the 26 NAM Lines. (1) Structural variants (SVs) were grouped based on their chromosomal position and size.(2) The first SV in each group was designated as the reference SV, and a 50 bp buffer (±25 bp from the start and end positions) was applied. Each remaining SV in the group was then compared to this reference (see Methods).(3–4) This process was iteratively repeated for all SVs until every SV was linked to a corresponding group.

**Figure S3.**
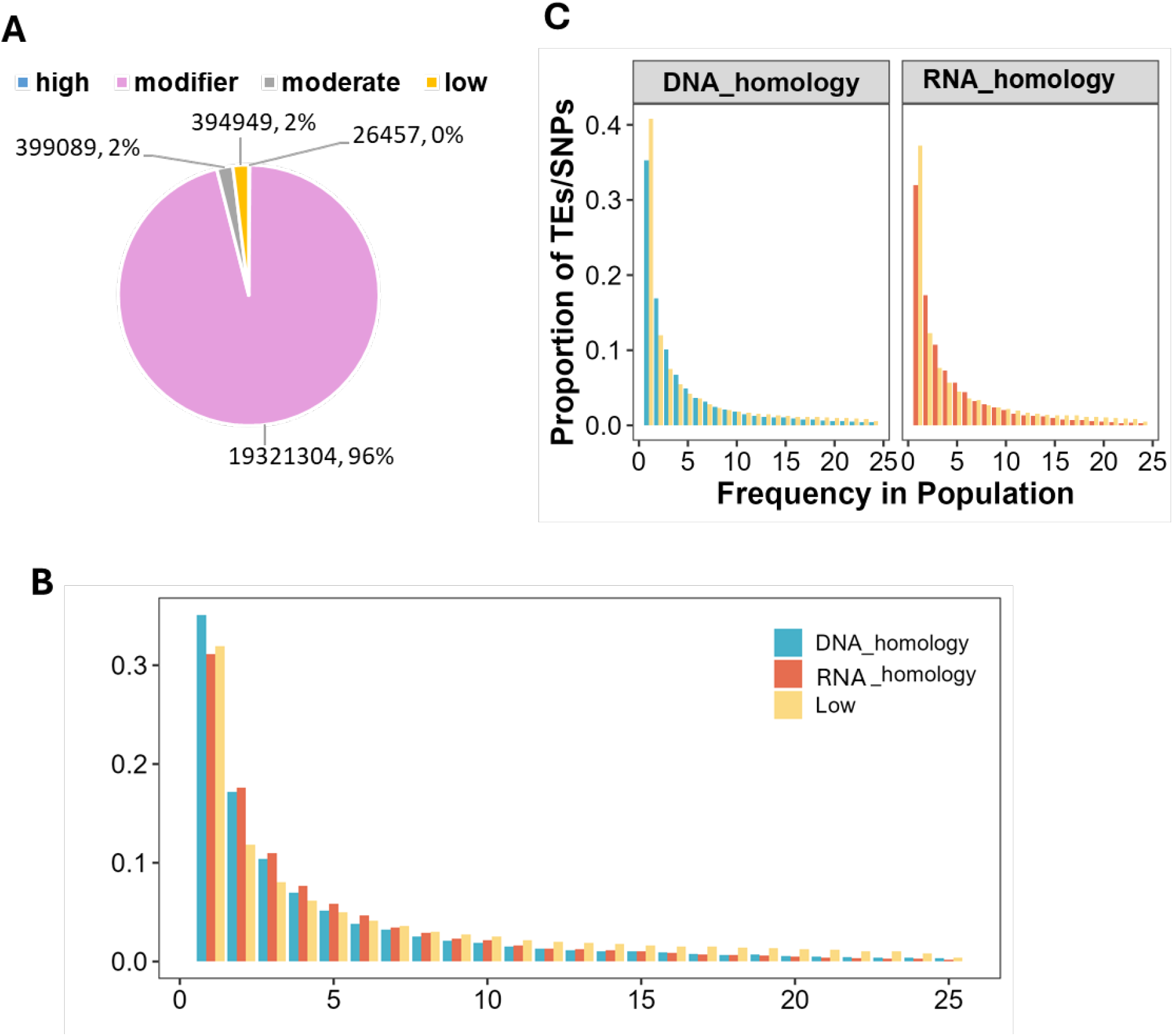
Site Frequency Spectrum (SFS) of DNA and RNA TEs identified by Sequence Homology. (A) The number/percentage of different single nucleotide polymorphisms (SNPs) groups.(B) Comparison of SFS distributions for DNA and RNA-homology TEs against Low SNP.(C) Comparison of SFS distributions for DNA and RNA-homology TEs with age-adjusted Low SNP distributions matched to each TE class.

**Figure S4.**
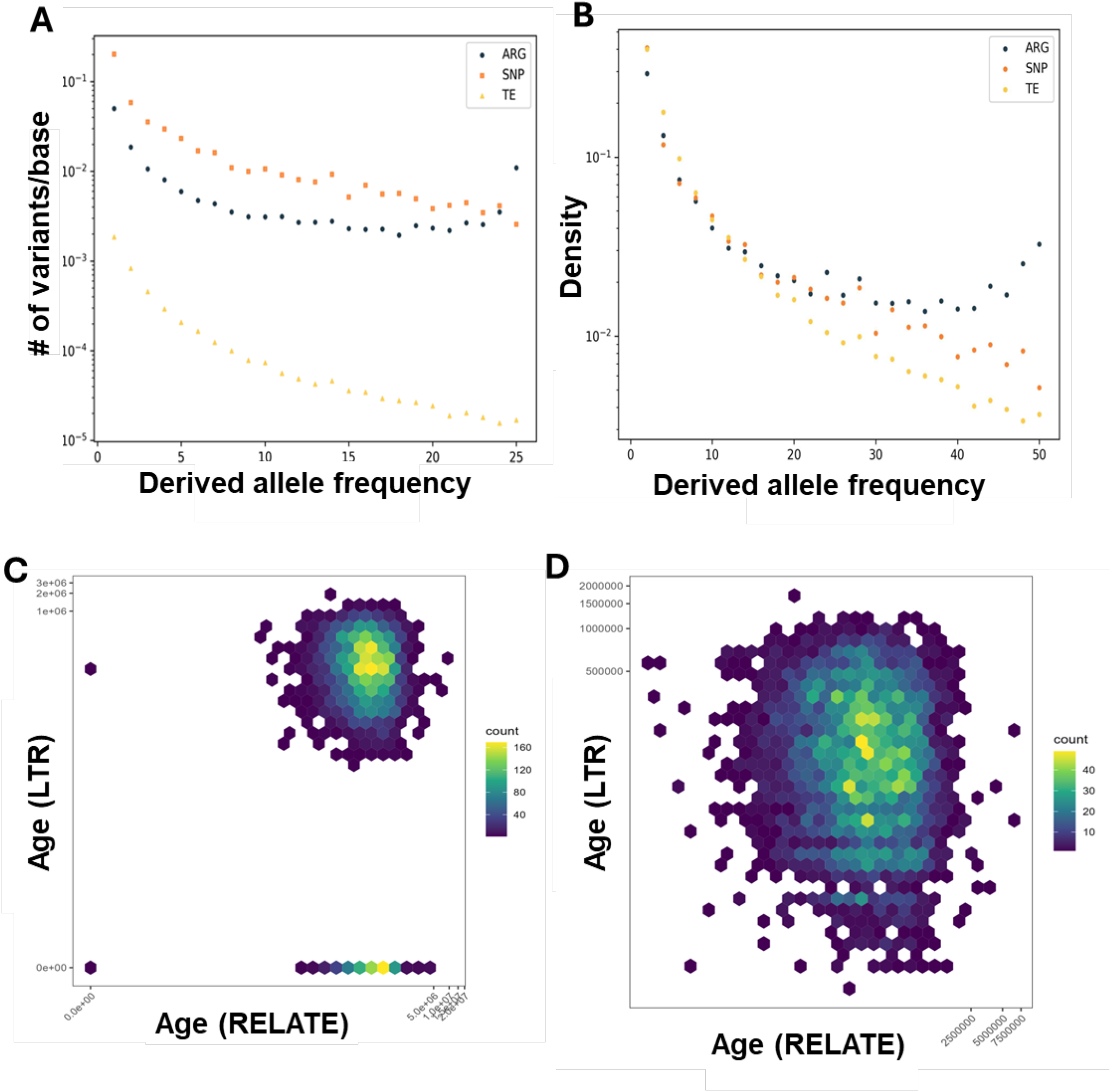
Validation of Age Estimation Accuracy Using RELATE. (A) Expected SFS from genealogical trees, also known as an ancestral recombination graph (ARG), from Relate and observed SFS from SNP and TE polymorphism data. (B) Normalized SFS. (C,D) Comparison of LTR retrotransposon insertion age with (C) and without (D) estimates of age 0 from RELATE with LTR sequence identity for insertions present in B73.

**Figure S5.**
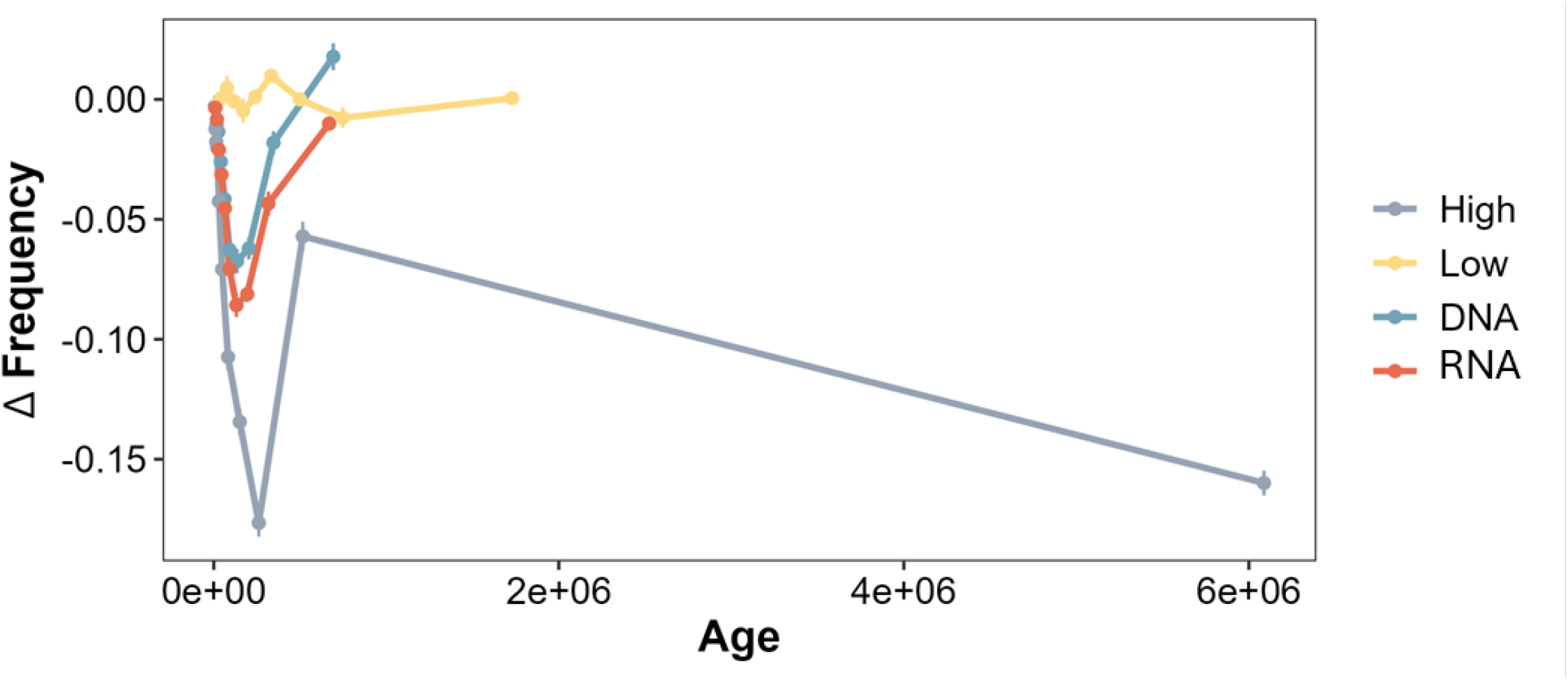
The Δ_*f*_ distribution of different SNPs (low and high) and TE (DNA and RNA) groups. Low and High SNPs are defined by their potential effects on genes.

**Figure S6.**
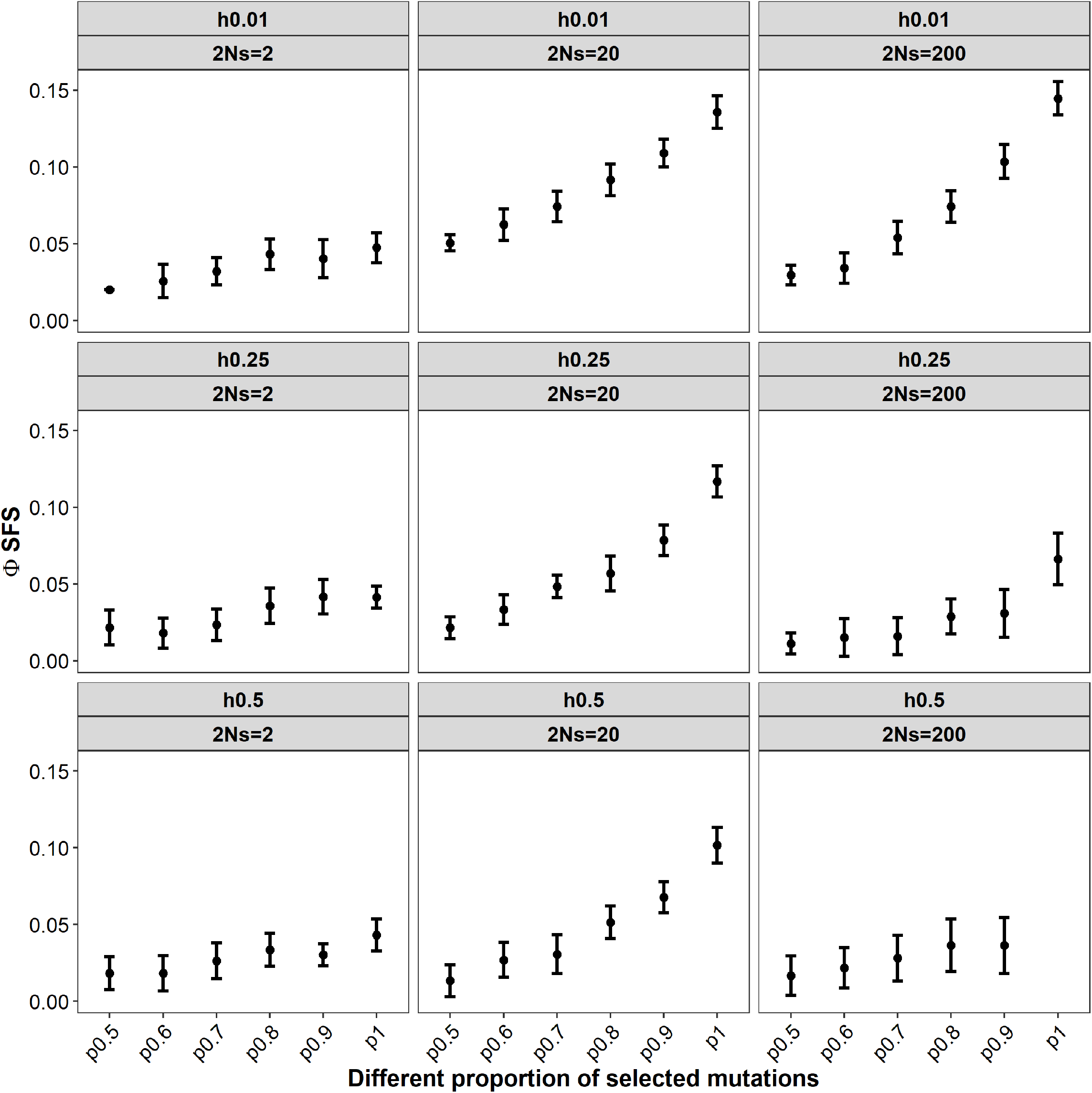
Simulated Φ_*SFS*_ values under varying evolutionary parameters. Φ_*SFS*_ was calculated from simulated datasets across combinations of dominance coefficients (*h* = 0.01, 0.25, 0.5), selection coefficients (2*Ns* = 2, 20, 200), and the proportion of mutations under selection (*p* = 0.5, 0.6, 0.7, 0.8, 0.9, 1).

**Figure S7.**
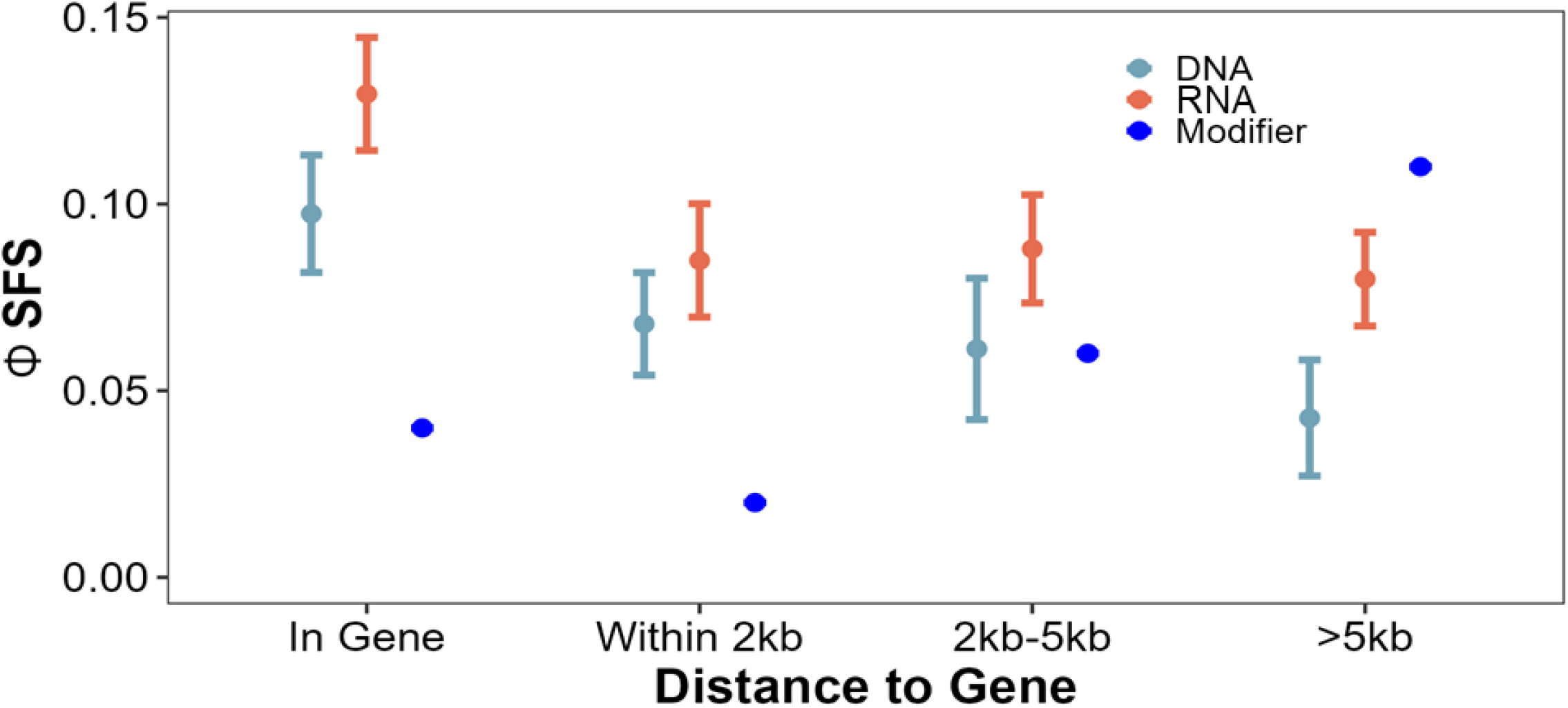
Comparison of Φ_*SFS*_ between modifier SNPs and TEs, including both DNA and RNA TEs, across different genomic locations.

**Figure S8.**
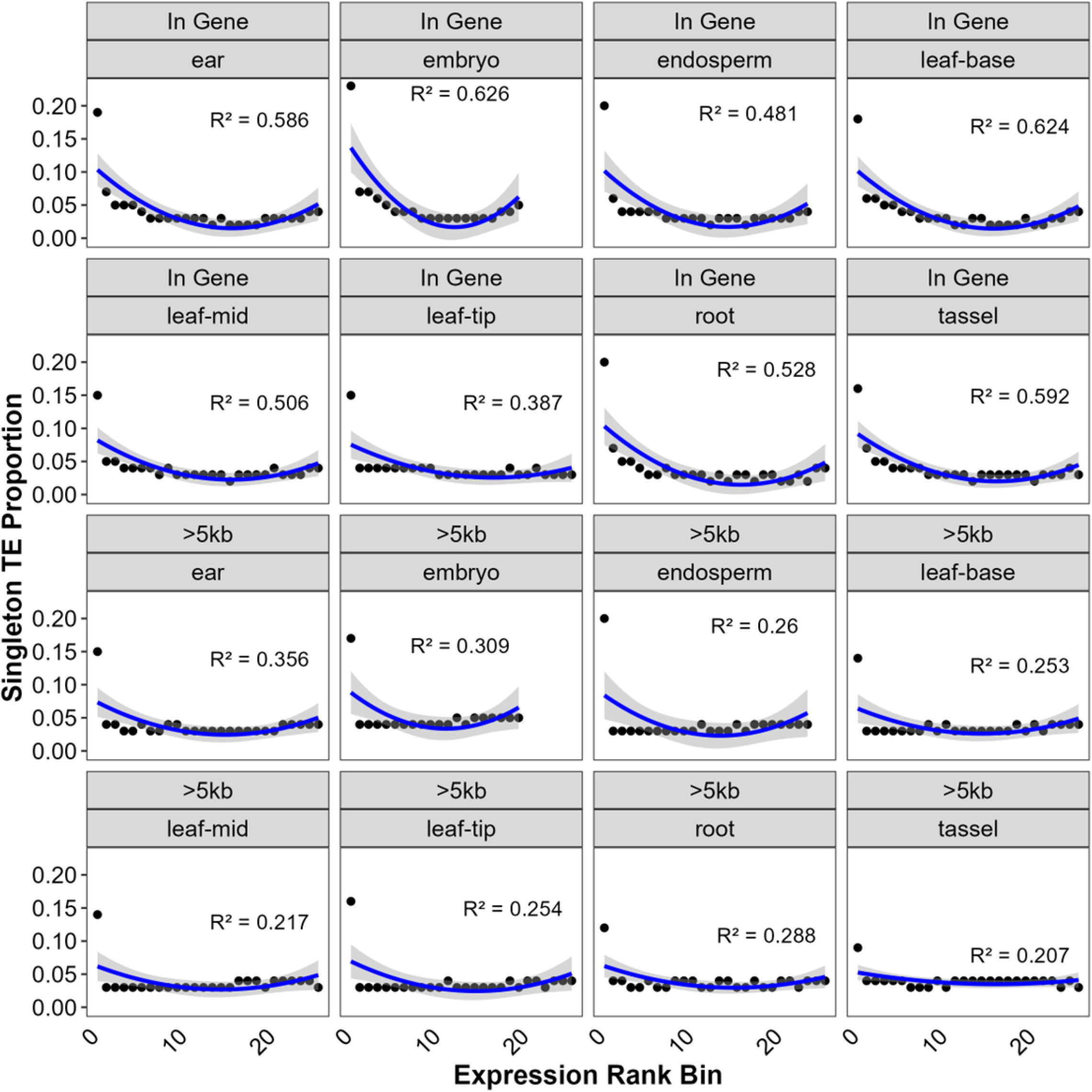
Relationship between singleton TE number and gene expression rank bin, separted by TE insertions into or > 5*Kb* distant from a gene in 8 different tissues.

**Figure S9.**
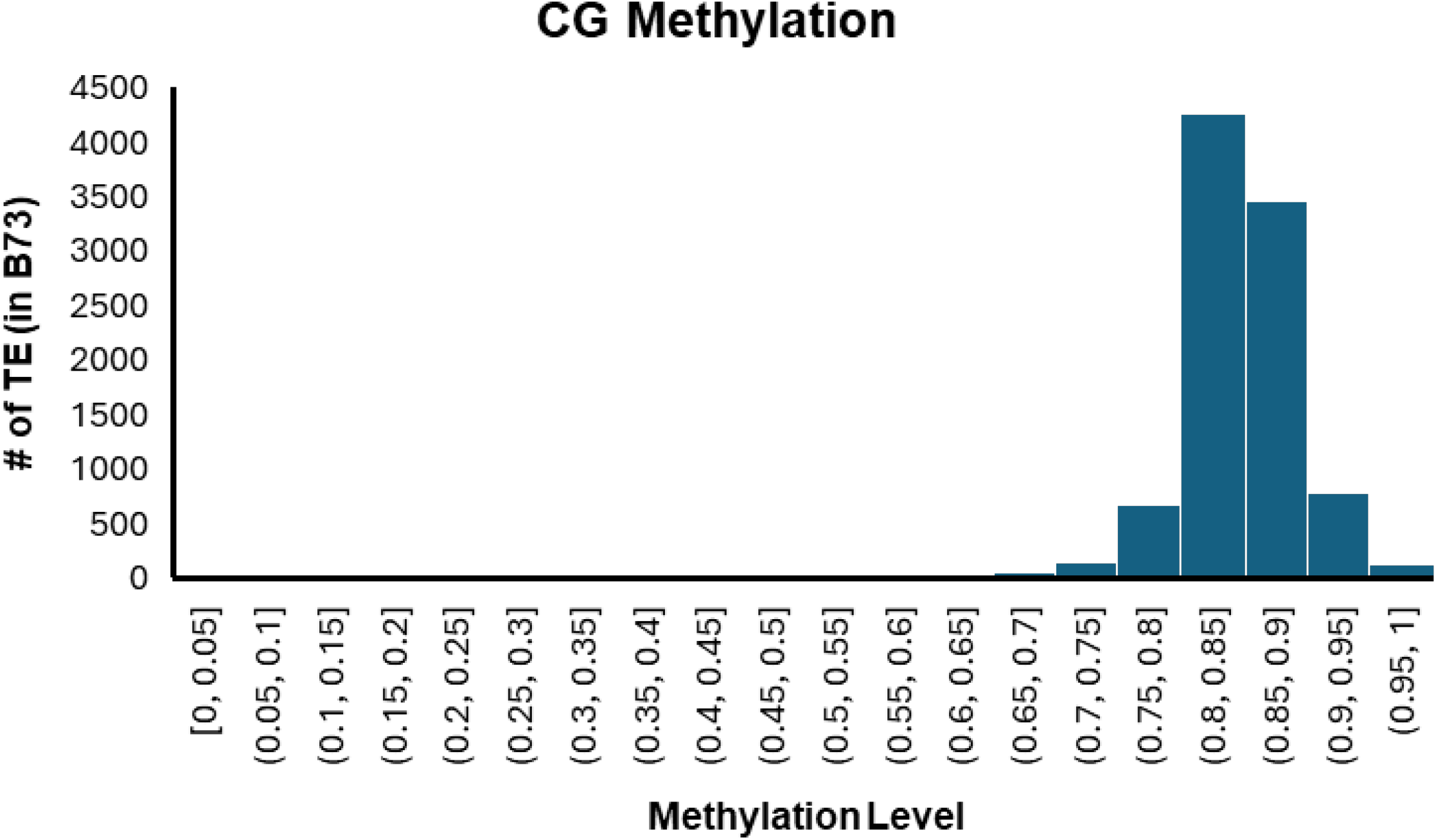
CG methylation patterns of TEs inserted in the B73 genome.

**Figure S10.**
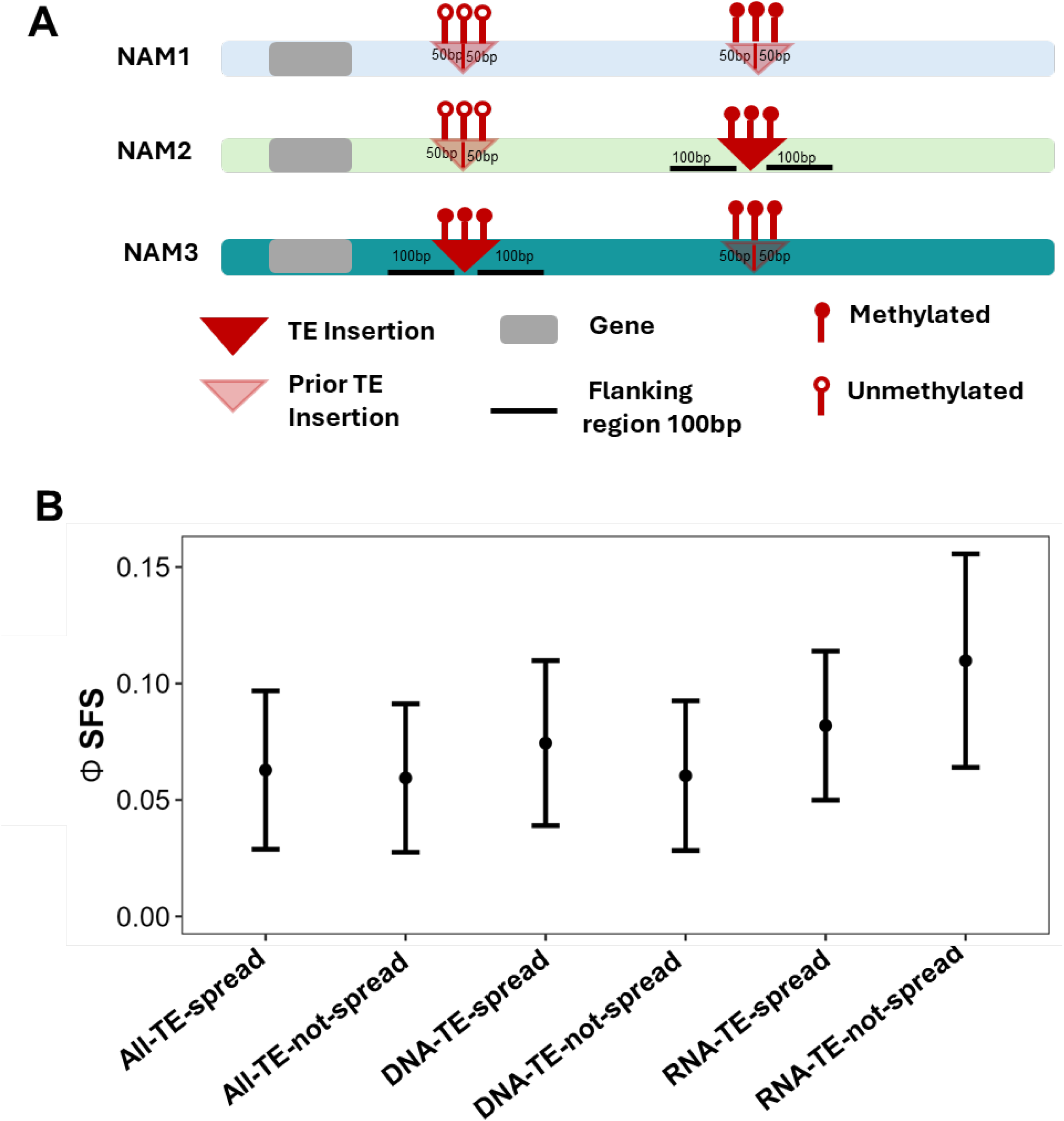
(A) Schematic illustrating the approach used to define genomic regions for assessing DNA methylation before and after TE insertion. The region prior to TE insertion was defined using homologous loci in NAM lines lacking the TE, while the region used to evaluate methylation spreading after TE insertion was defined as the 100 bp flanking sequence adjacent to the TE in lines where the TE is present.(B) The Φ_*S*_*FS* of TEs that do or do not exhibit methylation spreading to adjacent sequence. (Note: All-TE means DNA and RNA TE combined).

**Figure S11.**
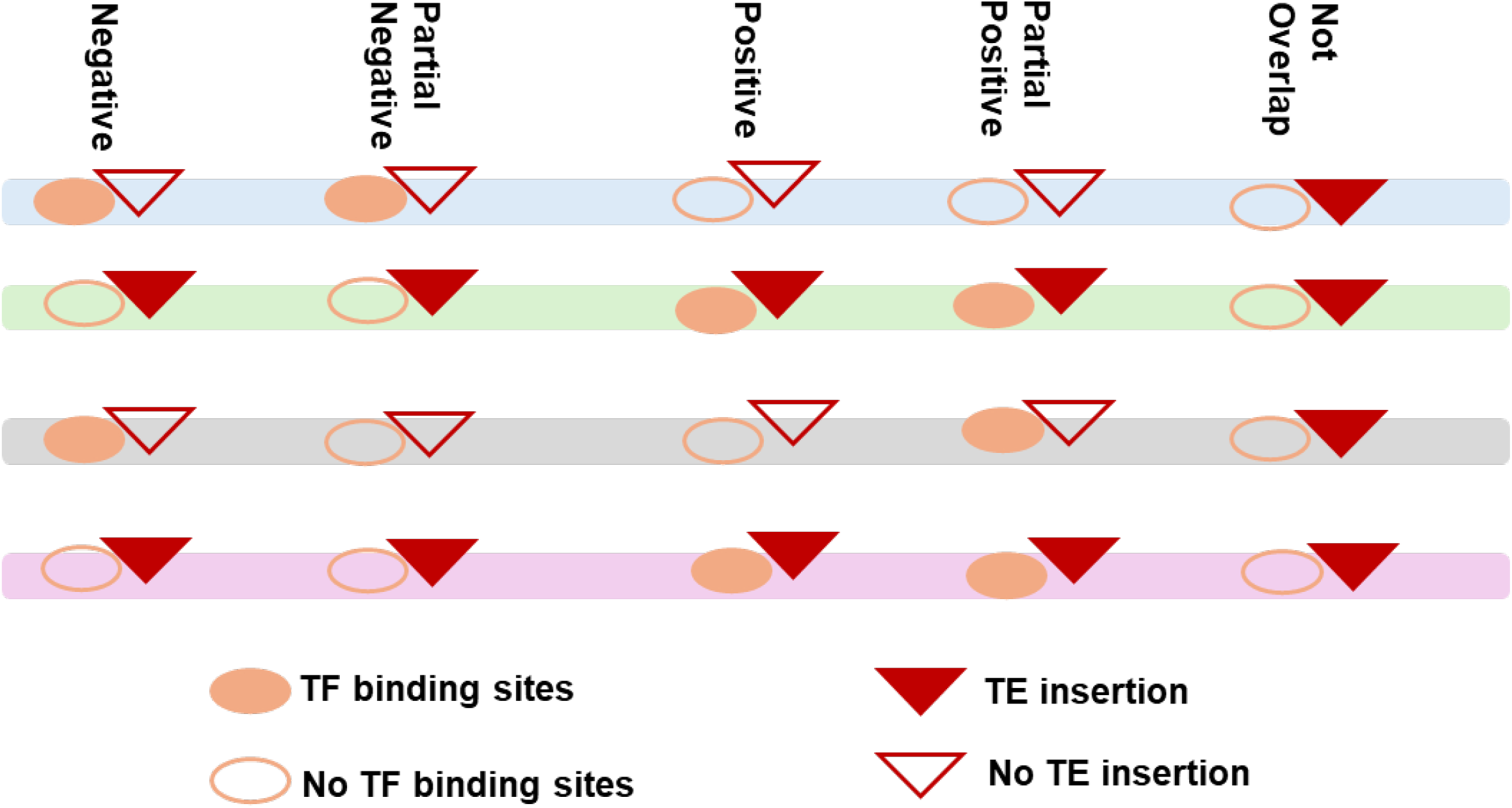
Schematic illustrating the different relationships between TEs and TF binding sites. TEs were categorized based on the presence or absence of TF binding in lines with and without the TE insertion.

**Figure S12.**
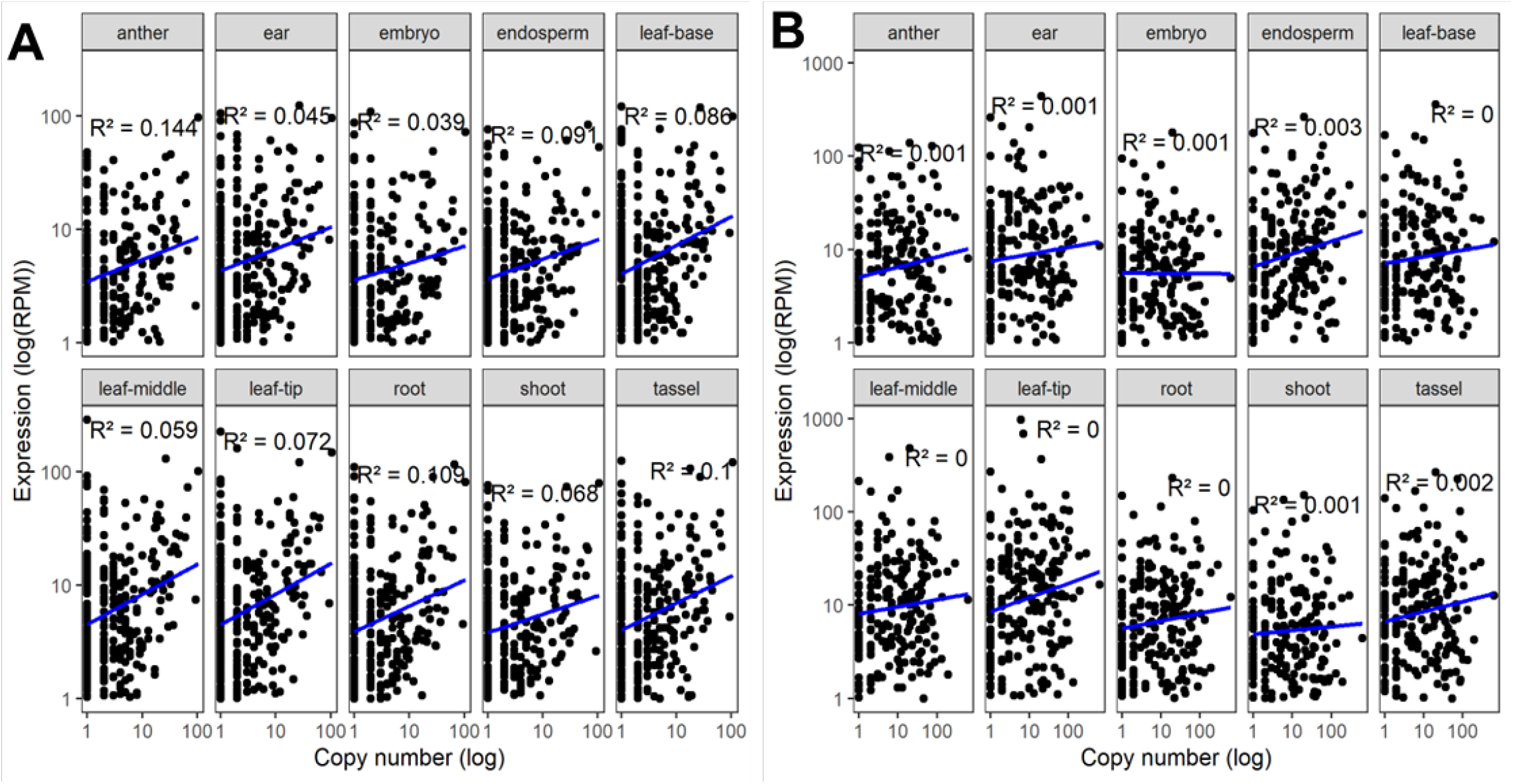
Correlation between (A) DNA and (B) RNA TE family level expression and copy number across ten tissues of B73.

**Figure S13.**
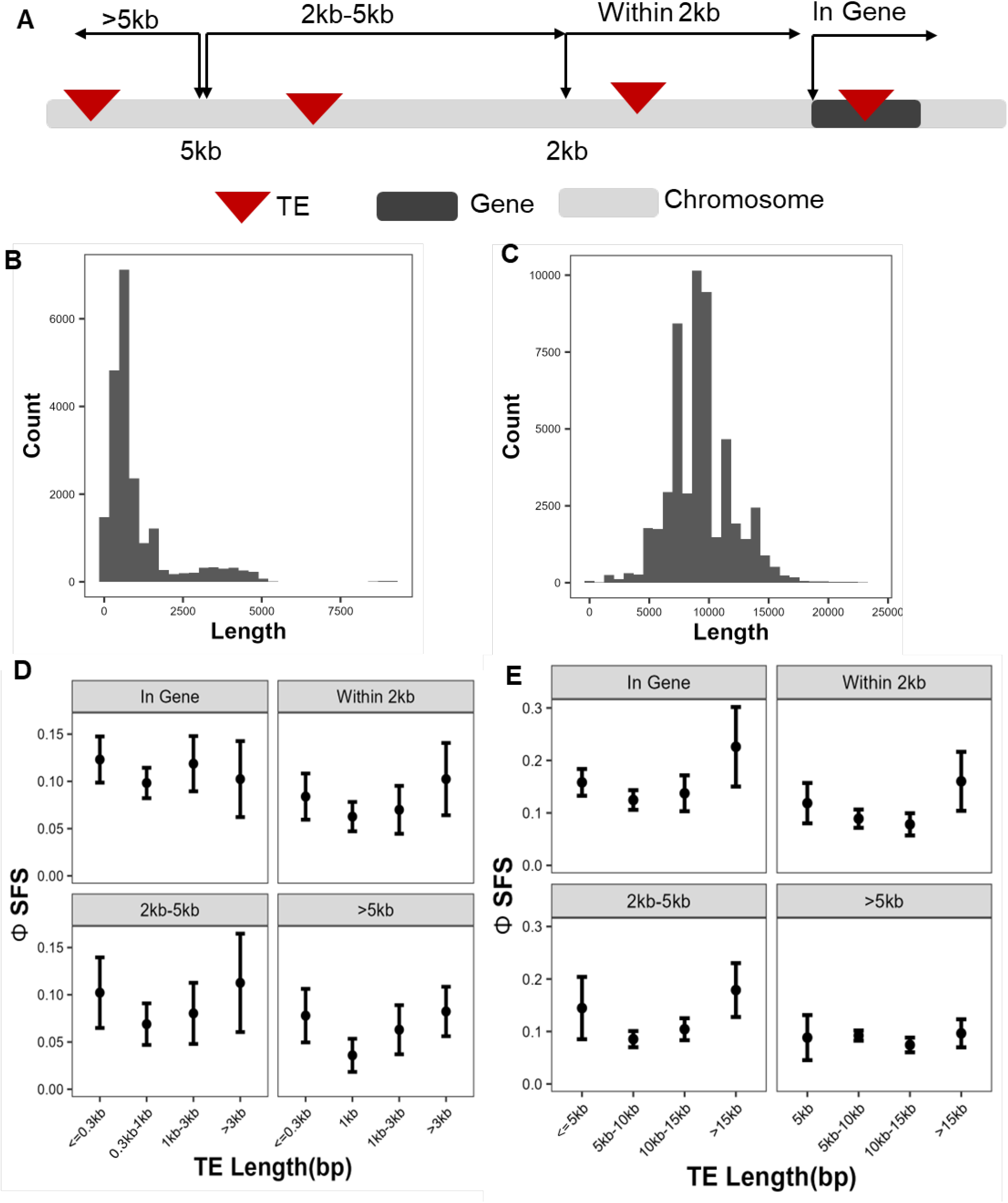
The effects of TE length on selection. (A) Schematic illustrating the logic used to define different genomic locations.(B–C) Length distribution of DNA and RNA TEs, respectively.(D–E) Φ_SFS_ values for TEs of different lengths across genomic locations for DNA and RNA TEs.

**Figure S14.**
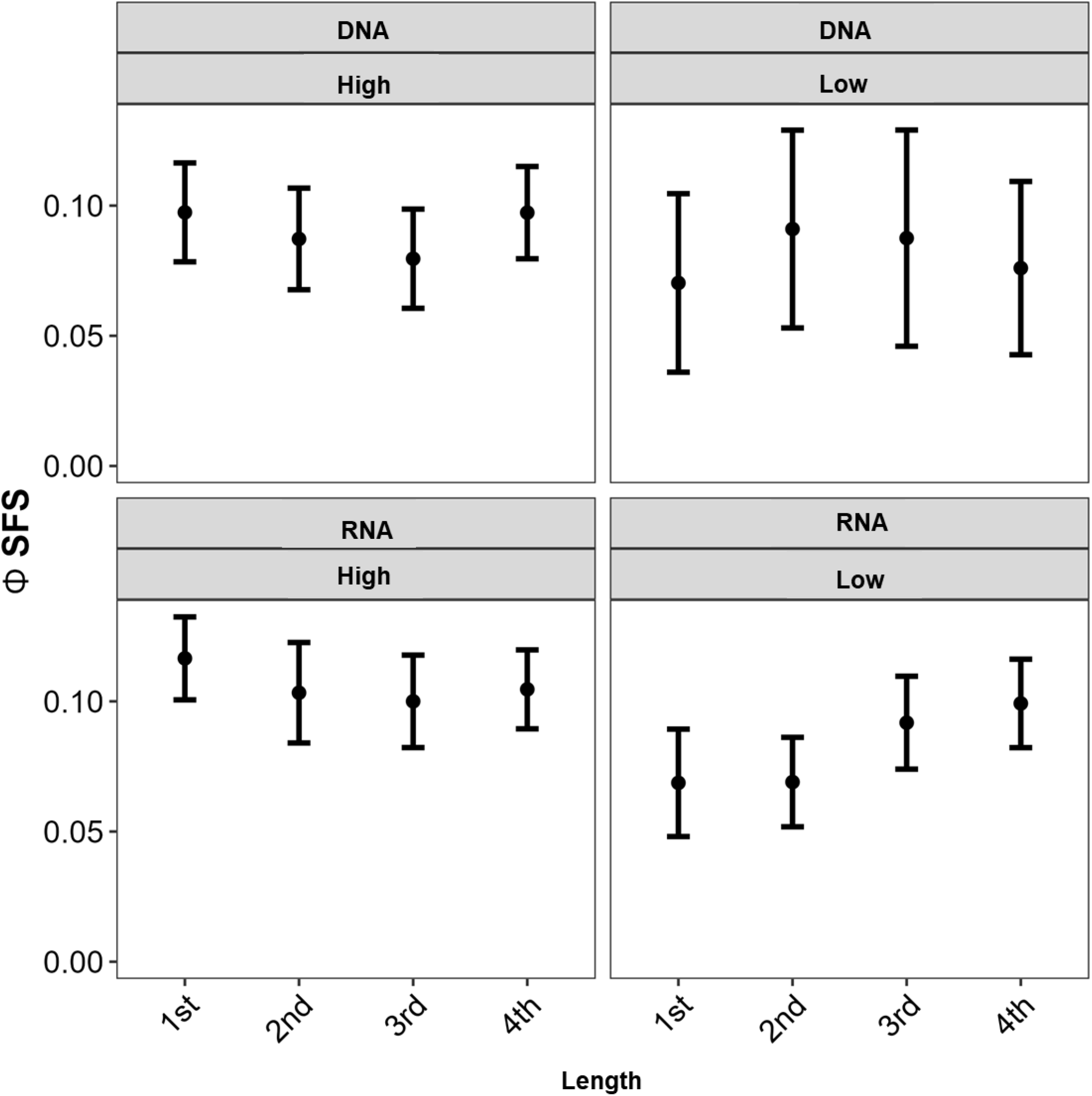
The selection of DNA and RNA TEs with different length (different quartiles) inserted into regions with high and low recombination rate.

**Figure S15.**
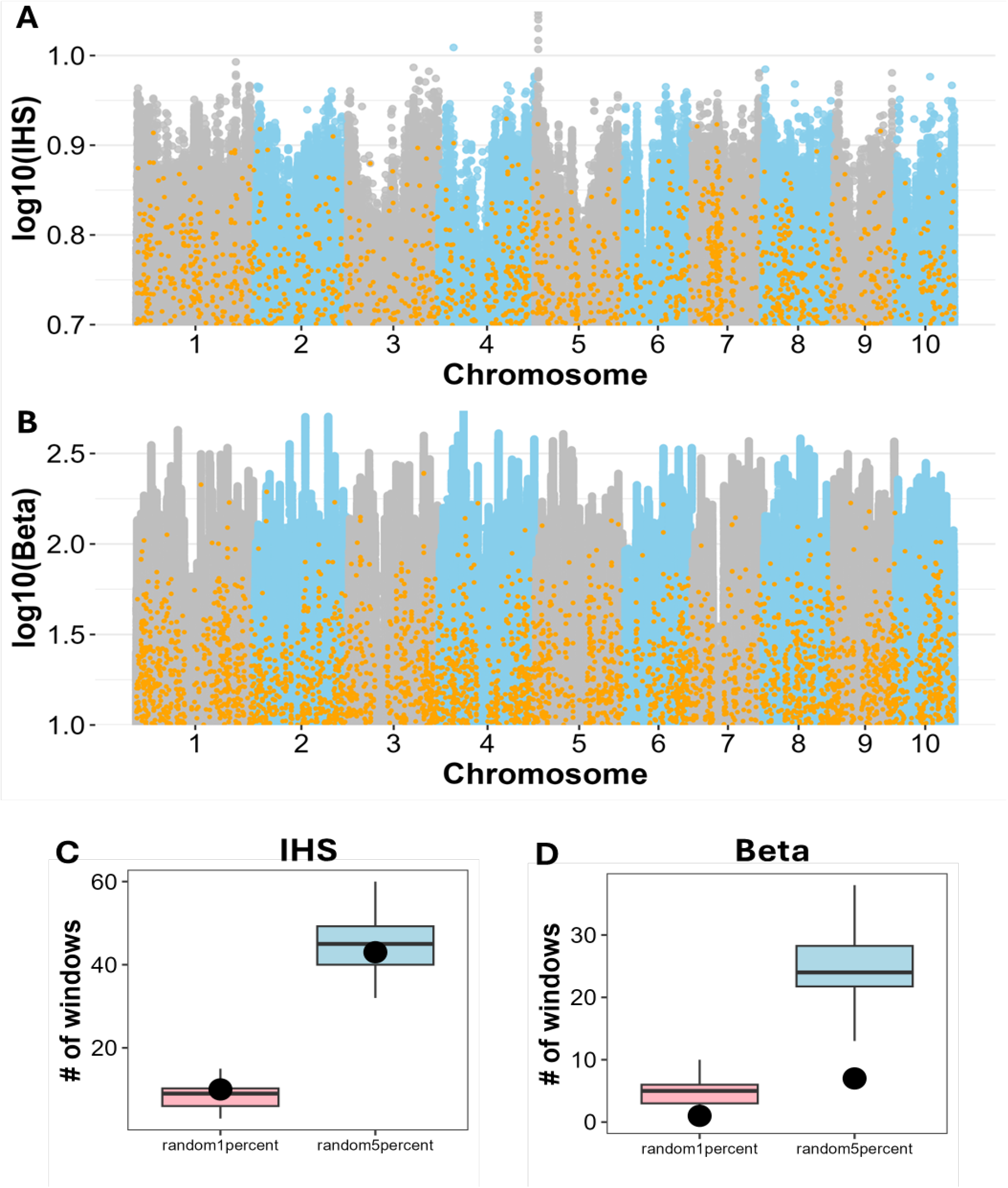
Genome-wide detection of TEs under positive and balancing selection. (A) Integrated Haplotype Score (iHS) of SNPs and TEs across the genome.(B) Beta score of SNPs and TEs across the genome. Number of windows where TEs have the highest (C) iHS and (D) Beta scores within randomly selected 1% and 5% of windows. (Notes, the black dots in (C) and (D) indicating the number of windows where TEs have the highest iHS and Beta score, respectively.

**Figure S16.**
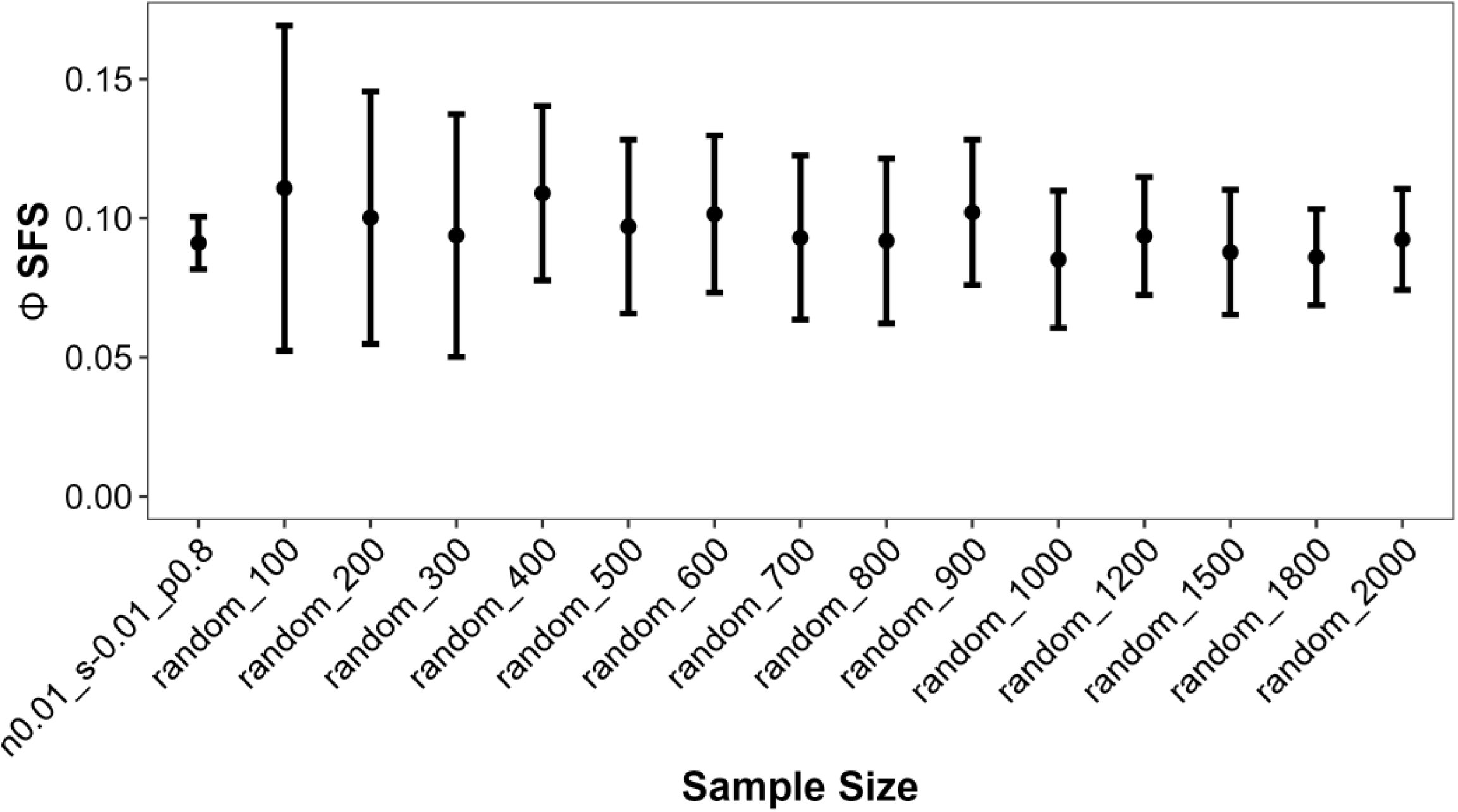
(A) Effect of sample size on the accuracy of the Φ_*SFS*_ method.

**Table S1.** Copy number of TEs across different superfamilies.

| Superfamily | Copy Number |
| --- | --- |
| RLC | 18690 |
| RLG | 26999 |
| RLX | 6710 |
| DTA | 11624 |
| DTC | 3108 |
| DTH | 4187 |
| DTM | 1377 |
| DTT | 319 |

**Table S2.** Copy number of TEs grouped by their relationship to transcription factor (TF) binding sites.

| TE-TF relationship | Copy Number |
| --- | --- |
| Negative | 378 |
| Partial Negative | 11519 |
| Positive | 117 |
| Partial Positive | 2554 |
| Not Overlap | 57150 |

## References

Aminetzach, Y. T., J. M. Macpherson, and D. A. Petrov, 2005 Pesticide resistance via transposition-mediated adaptive gene truncation in drosophila. Science 309: 764–767.

Anderson, S. N., G. J. Zynda, J. Song, Z. Han, M. W. Vaughn, et al., 2018 Subtle perturbations of the maize methylome reveal genes and transposons silenced by chromomethylase or rna-directed dna methylation pathways. G3: Genes, Genomes, Genetics 8: 1921–1932.

Badge, R. M. and J. F. Brookfield, 1997 The role of host factors in the population dynamics of selfish transposable elements. Journal of theoretical biology 187: 261–271.

Baduel, P., B. Leduque, A. Ignace, I. Gy, J. Gil Jr, et al., 2021 Genetic and environmental modulation of transposition shapes the evolutionary potential of arabidopsis thaliana. Genome biology 22: 138.

Baumdicker, F., G. Bisschop, D. Goldstein, G. Gower, A. P. Ragsdale, et al., 2022 Efficient ancestry and mutation simulation with msprime 1.0. Genetics 220: iyab229.

Beissinger, T., 2014 Genwin: Spline based window boundaries for genomic analyses.

Beissinger, T. M., L. Wang, K. Crosby, A. Durvasula, M. B. Hufford, et al., 2016 Recent demography drives changes in linked selection across the maize genome. Nature plants 2: 1–7.

Betancourt, A. J., K. H.-C. Wei, Y. Huang, and Y. C. G. Lee, 2024 Causes and consequences of varying transposable element activity: An evolutionary perspective. Annual review of genomics and human genetics 25: 1–25.

Bhattacharyya, M. K., A. M. Smith, T. N. Ellis, C. Hedley, and C. Martin, 1990 The wrinkled-seed character of pea described by mendel is caused by a transposon-like insertion in a gene encoding starch-branching enzyme. Cell 60: 115–122.

Bourque, G., K. H. Burns, M. Gehring, V. Gorbunova, A. Seluanov, et al., 2018 Ten things you should know about transposable elements. Genome biology 19: 199.

Bubb, K. L., M. O. Hamm, T. W. Tullius, J. K. Min, B. Ramirez-Corona, et al., 2025 The regulatory potential of transposable elements in maize. Nature Plants pp. 1–12.

Butelli, E., C. Licciardello, Y. Zhang, J. Liu, S. Mackay, et al., 2012 Retrotransposons control fruit-specific, cold-dependent accumulation of anthocyanins in blood oranges. The Plant Cell 24: 1242–1255.

Casillas, S., A. Barbadilla, and C. M. Bergman, 2007 Purifying selection maintains highly conserved noncoding sequences in drosophila. Molecular biology and evolution 24: 2222–2234.

Charlesworth, B., 2013 Stabilizing selection, purifying selection, and mutational bias in finite populations. Genetics 194: 955–971.

Choi, J. Y. and M. D. Purugganan, 2018 Evolutionary epigenomics of retrotransposon-mediated methylation spreading in rice. Molecular Biology and Evolution 35: 365–382.

Cingolani, P., A. Platts, M. Coon, T. Nguyen, L. Wang, et al., 2012 A program for annotating and predicting the effects of single nucleotide polymorphisms, snpeff: Snps in the genome of drosophila melanogaster strain w1118; iso-2; iso-3. Fly 6: 80–92.

Clark, R. M., S. Tavaré, and J. Doebley, 2005 Estimating a nucleotide substitution rate for maize from polymorphism at a major domestication locus. Molecular biology and evolution 22: 2304–2312.

Engelhorn, J., S. J. Snodgrass, A. Kok, A. S. Seetharam, M. Schneider, et al., 2024 Genetic variation at transcription factor binding sites largely explains phenotypic heritability in maize. bioRxiv 10: 08–551183.

Haller, B. C. and P. W. Messer, 2019a Slim 3: forward genetic simulations beyond the wright–fisher model. Molecular biology and evolution 36: 632–637.

Haller, B. C. and P. W. Messer, 2019b Slim 3: Forward genetic simulations beyond the wright–fisher model. Molecular Biology and Evolution 36: 632–637.

Hénault, M., S. Marsit, G. Charron, and C. R. Landry, 2020 The effect of hybridization on transposable element accumulation in an undomesticated fungal species. Elife 9: e60474.

Hof, A. E. v., P. Campagne, D. J. Rigden, C. J. Yung, J. Lingley, et al., 2016 The industrial melanism mutation in british peppered moths is a transposable element. Nature 534: 102–105.

Hollister, J. D. and B. S. Gaut, 2009 Epigenetic silencing of transposable elements: a trade-off between reduced transposition and deleterious effects on neighboring gene expression. Genome research 19: 1419–1428.

Horvath, R., M. Menon, M. Stitzer, and J. Ross-Ibarra, 2022 Controlling for variable transposition rate with an age-adjusted site frequency spectrum. Genome Biology and Evolution 14: evac016.

Horvath, R., N. Minadakis, Y. Bourgeois, and A. C. Roulin, 2024 The evolution of transposable elements in Brachypodium distachyon is governed by purifying selection, while neutral and adaptive processes play a minor role. eLife 12: RP93284.

Horvath, R. and T. Slotte, 2017 The role of small rna-based epigenetic silencing for purifying selection on transposable elements in capsella grandiflora. Genome Biology and Evolution 9: 2911–2920.

Huang, C., H. Sun, D. Xu, Q. Chen, Y. Liang, et al., 2018 Zmcct9 enhances maize adaptation to higher latitudes. Proceedings of the National Academy of Sciences 115: E334–E341.

Huang, Y. and Y. C. G. Lee, 2024 Blessing or curse: how the epigenetic resolution of host-transposable element conflicts shapes their evolutionary dynamics. Proceedings of the Royal Society B 291: 20232775.

Hufford, M. B., A. S. Seetharam, M. R. Woodhouse, K. M. Chougule, S. Ou, et al., 2021 De novo assembly, annotation, and comparative analysis of 26 diverse maize genomes. Science 373: 655–662.

Jiang, J., Y.-C. Xu, Z.-Q. Zhang, J.-F. Chen, X.-M. Niu, et al., 2024 Forces driving transposable element load variation during arabidopsis range expansion. The Plant Cell 36: 840–862.

Kelleher, J., A. M. Etheridge, and G. McVean, 2016 Efficient coalescent simulation and genealogical analysis for large sample sizes. PLoS computational biology 12: e1004842.

Kent, T. V., J. Uzunović, and S. I. Wright, 2017 Coevolution between transposable elements and recombination. Philosophical Transactions of the Royal Society B: Biological Sciences 372: 20160458.

Kobayashi, S., N. Goto-Yamamoto, and H. Hirochika, 2004 Retrotransposon-induced mutations in grape skin color. Science 304: 982–982.

Kofler, R., A. J. Betancourt, and C. Schlötterer, 2012 Sequencing of pooled dna samples (pool-seq) uncovers complex dynamics of transposable element insertions in drosophila melanogaster. PLoS genetics 8: e1002487.

Kremling, K. A., S.-Y. Chen, M.-H. Su, N. K. Lepak, M. C. Romay, et al., 2018 Dysregulation of expression correlates with rare-allele burden and fitness loss in maize. Nature 555: 520–523.

Li, X., X. Dai, H. He, Y. Lv, L. Yang, et al., 2024 A pante map highlights transposable elements underlying domestication and agronomic traits in asian rice. National Science Review 11: nwae188.

Lisch, D., 2009 Epigenetic regulation of transposable elements in plants. Annual review of plant biology 60: 43–66.

Lisch, D., 2013 How important are transposons for plant evolution? Nature reviews genetics 14: 49–61.

Liu, B. and M. Zhao, 2023 How transposable elements are recognized and epigenetically silenced in plants? Current Opinion in Plant Biology 75: 102428.

Makarevitch, I., A. J. Waters, P. T. West, M. Stitzer, C. N. Hirsch, et al., 2015 Transposable elements contribute to activation of maize genes in response to abiotic stress. PLoS genetics 11: e1004915.

McClintock, B., 1950 The origin and behavior of mutable loci in maize. Proceedings of the National Academy of Sciences 36: 344–355.

McKenna, A., M. Hanna, E. Banks, A. Sivachenko, K. Cibulskis, et al., 2010 The genome analysis toolkit: a mapreduce framework for analyzing next-generation dna sequencing data. Genome research 20: 1297–1303.

McMullen, M. D., S. Kresovich, H. S. Villeda, P. Bradbury, H. Li, et al., 2009 Genetic properties of the maize nested association mapping population. Science 325: 737–740.

Menard, C., N. Catlin, A. Platts, Y. Qiu, E. Roback, et al., 2025 Swif-te: identifying novel transposable element insertions from short read data. bioRxiv pp. 2025–07.

Miles, A., pyup.io bot, M. F. Rodrigues, P. Ralph, J. Kelleher, et al., 2024 cggh/scikit-allel: v1.3.13.

Munasinghe, M., A. Read, M. C. Stitzer, B. Song, C. C. Menard, et al., 2023a Combined analysis of transposable elements and structural variation in maize genomes reveals genome contraction outpaces expansion. PLoS genetics 19: e1011086.

Munasinghe, M., N. Springer, and Y. Brandvain, 2023b Critical role of insertion preference for invasion trajectory of transposons. Evolution 77: 2173–2185.

Naito, K., E. Cho, G. Yang, M. A. Campbell, K. Yano, et al., 2006 Dramatic amplification of a rice transposable element during recent domestication. Proceedings of the National Academy of Sciences 103: 17620–17625.

Neafsey, D. E., J. P. Blumenstiel, and D. L. Hartl, 2004 Different regulatory mechanisms underlie similar transposable element profiles in pufferfish and fruitflies. Molecular Biology and Evolution 21: 2310–2318.

Nielsen, R., 2005 Molecular signatures of natural selection. Annu. Rev. Genet. 39: 197–218.

Noshay, J. M., S. N. Anderson, P. Zhou, L. Ji, W. Ricci, et al., 2019 Monitoring the interplay between transposable element families and dna methylation in maize. PLoS Genetics 15: e1008291.

Noshay, J. M., A. P. Marand, S. N. Anderson, P. Zhou, M. K. Mejia Guerra, et al., 2021 Assessing the regulatory potential of transposable elements using chromatin accessibility profiles of maize transposons. Genetics 217: 1–13.

O’Neill, R. J. W., M. J. O’Neill, and J. A. M. Graves, 1998 Undermethylation associated with retroelement activation and chromosome remodelling in an interspecific mammalian hybrid. Nature 393: 68–72.

Ong-Abdullah, M., J. M. Ordway, N. Jiang, S.-E. Ooi, S.-Y. Kok, et al., 2015 Loss of karma transposon methylation underlies the mantled somaclonal variant of oil palm. Nature 525: 533–537.

Petrov, D. A., A.-S. Fiston-Lavier, M. Lipatov, K. Lenkov, and J. González, 2011 Population genomics of transposable elements in drosophila melanogaster. Molecular biology and evolution 28: 1633–1644.

Preston, B. D., 1996 Error-prone retrotransposition: rime of the ancient mutators. Proceedings of the National Academy of Sciences 93: 7427–7431.

Quinlan, A. R. and I. M. Hall, 2010 Bedtools: a flexible suite of utilities for comparing genomic features. Bioinformatics 26: 841–842.

Ralph, P., K. Thornton, and J. Kelleher, 2020 Efficiently summarizing relationships in large samples: A general duality between statistics of genealogies and genomes. Genetics 215: 779–797.

SanMiguel, P., B. S. Gaut, A. Tikhonov, Y. Nakajima, and J. L. Bennetzen, 1998 The paleontology of intergene retrotransposons of maize. Nature genetics 20: 43–45.

Siewert, K. M. and B. F. Voight, 2017 Detecting long-term balancing selection using allele frequency correlation. Molecular biology and evolution 34: 2996–3005.

Sigman, M. J. and R. K. Slotkin, 2016 The first rule of plant transposable element silencing: location, location, location. The Plant Cell 28: 304–313.

Song, M. and S. Boissinot, 2007 Selection against line-1 retro-transposons results principally from their ability to mediate ectopic recombination. Gene 390: 206–213.

Speidel, L., M. Forest, S. Shi, and S. R. Myers, 2019 A method for genome-wide genealogy estimation for thousands of samples. Nature genetics 51: 1321–1329.

Stitzer, M. C., S. N. Anderson, N. M. Springer, and J. Ross-Ibarra, 2021 The genomic ecosystem of transposable elements in maize. PLoS genetics 17: e1009768.

Stitzer, M. C., M. B. Khaipho-Burch, A. I. Hudson, B. Song, J. A. Valdez-Franco, et al., 2023 Transposable element abundance subtly contributes to lower fitness in maize. bioRxiv pp. 2023–09.

Studer, A., Q. Zhao, J. Ross-Ibarra, and J. Doebley, 2011 Identification of a functional transposon insertion in the maize domestication gene tb1. Nature genetics 43: 1160–1163.

Szpiech, Z. A. and R. D. Hernandez, 2014 selscan: an efficient multithreaded program to perform ehh-based scans for positive selection. Molecular biology and evolution 31: 2824–2827.

Torkamanzehi, A., C. Moran, and F. W. Nicholas, 1992 P element transposition contributes substantial new variation for a quantitative trait in drosophila melanogaster. Genetics 131: 73–78.

Ungerer, M. C., S. C. Strakosh, and K. M. Stimpson, 2009 Proliferation of ty3/gypsy-like retrotransposons in hybrid sunflower taxa inferred from phylogenetic data. Bmc Biology 7: 40.

Uzunović, J., E. B. Josephs, J. R. Stinchcombe, and S. I. Wright, 2019 Transposable elements are important contributors to standing variation in gene expression in capsella grandiflora. Molecular biology and evolution 36: 1734–1745.

Voight, B. F., S. Kudaravalli, X. Wen, and J. K. Pritchard, 2006 A map of recent positive selection in the human genome. PLoS biology 4: e72.

Wei, K. H.-C., D. Mai, K. Chatla, and D. Bachtrog, 2022 Dynamics and impacts of transposable element proliferation in the drosophila nasuta species group radiation. Molecular Biology and Evolution 39: msac080.

Wicker, T., F. Sabot, A. Hua-Van, J. L. Bennetzen, P. Capy, et al., 2007 A unified classification system for eukaryotic transposable elements. Nature reviews genetics 8: 973–982.

Wong, Y., A. Ignatieva, J. Koskela, G. Gorjanc, A. W. Wohns, et al., 2024 A general and efficient representation of ancestral recombination graphs. Genetics 228: iyae100.

Xiao, H., N. Jiang, E. Schaffner, E. J. Stockinger, and E. van der Knaap, 2008 A retrotransposon-mediated gene duplication underlies morphological variation of tomato fruit. science 319: 1527–1530.

Yang, Q., Z. Li, W. Li, L. Ku, C. Wang, et al., 2013 Cacta-like transposable element in zmcct attenuated photoperiod sensitivity and accelerated the postdomestication spread of maize. Proceedings of the National Academy of Sciences 110: 16969–16974.

Zeng, Y., R. K. Dawe, and J. I. Gent, 2023 Natural methylation epialleles correlate with gene expression in maize. Genetics 225: iyad146.

Zhao, J., I. Akinsanmi, D. Arafat, T. Cradick, C. M. Lee, et al., 2016 A burden of rare variants associated with extremes of gene expression in human peripheral blood. The American Journal of Human Genetics 98: 299–309.

Zhao, M., B. Zhang, D. Lisch, and J. Ma, 2017 Patterns and consequences of subgenome differentiation provide insights into the nature of paleopolyploidy in plants. The Plant Cell 29: 2974–2994.

